# Balancing off-target and on-target considerations for optimized Cas9 CRISPR knockout library design

**DOI:** 10.1101/2025.08.26.672375

**Authors:** Laura M Drepanos, Smriti Srikanth, Eleanor G Kaplan, Spencer T Shah, Berta Escude Velasco, Sarra Merzouk, John G Doench

## Abstract

The continued development of high-dimensional CRISPR screen readouts, such as single-cell RNA sequencing and high-content imaging, necessitates compact libraries to enable functional interrogation at genome scale. Improved genome annotations yield library deprecation over time, further motivating an updated genome-wide design effort. Recently, we have developed an enhanced model, Rule Set 3, which leveraged an expansive training set and feature space to predict guide efficacy. However, the benefit of such advances to library design is limited by current approaches to balance predictions of on-target activity with off-target considerations. Here we present a guide selection strategy that identifies guides with sufficient off-target activity to justify omission from the library, thus avoiding the unnecessary exclusion of active guides. We pair this model with strategic design choices to create Jacquere, an updated, optimized, and validated Cas9 CRISPR knockout (CRISPRko) genome-wide library for the human genome.

## INTRODUCTION

CRISPR screens, which reveal the phenotype resulting from a genetic perturbation, are an effective approach to probe gene function ^1–3^. Since guide RNAs (guides) may fail to perturb the targeted gene or yield a phenotype independent of the gene’s function due to activity at off-target sites, herein referred to as guide promiscuity, genes must be targeted by numerous guides to reliably infer their function from the observed results. However, the large scale and cost of screens, especially among approaches that associate perturbations with multidimensional phenotypes such as Perturb-seq and Optical Pooled Screens, limits the depth at which each gene can be targeted ^4–8^. Such advances therefore necessitate careful selection of guides that are effective at the target site with minimal promiscuity.

Numerous tools support the design of screening libraries. For individual guide assessment, the command line tools FlashFry and CALITAS and web tools GT-Scan, Cas-OFFinder, and CRISPRitz enumerate potential off-target sites ^9–13^. The CRISPOR web tool evaluates guides for both on-target and off-target properties, but is limited in its capabilities for batch guide design ^14^. Alternatively, the web tool CRISPick (developed by our group) as well as Guidescan2 enable ad-hoc CRISPR screening library design for the Cas enzyme, modality, and target genome of interest, scoring candidate guides for user-specified targets in batch with respect to on-target and off-target properties and balancing such criteria to select the user-defined number of guides for each gene ^15,16^. The CRISPick guide design tool updates continuously to feature current NCBI and Ensembl genome annotations and iterative improvements to guide design, which are reflected in genome scale guide selections available for download.

Further, numerous genome-wide guide libraries have been pre-assembled for widespread usage. Among libraries designed for knockout screens with Cas9 from *Streptococcus pyogenes* (hereafter simply Cas9), GeCKO v2 reflects an early effort to select guides with minimal off-target activity^17^. In contrast, the Avana library prioritizes guides with substantial efficacy as predicted by the 2014 Rule Set 1 model of on-target activity and generally avoids guides with perfect off-target matches^18^. This library has been extensively screened by the Dependency Map (DepMap) project at the Broad Institute ^19^. Likewise, the Sanger Institute’s Project Score effort utilizes a genome-wide library that reflects both on-target and off-target considerations ^20,21^. The Brunello library incorporated improved guide activity predictions with Rule Set 2 and the development of the Cutting Frequency Determination (CFD) score to predict activity rates at off-target sites^15^. The TKOv3 library, released shortly afterwards, similarly incorporates predictions of on-target and off-target activity in guide designs^22^. Acknowledging the screening limitations introduced by in-frame mutations, which comprise approximately 20% of Cas9 CRISPRko edits, the Vienna Bioactivity CRISPR (VBC) score incorporates Rule Set 2 predictions of guide efficacy with predictions of mutational outcomes and amino acid-level consequences for library design^23^. Recently, more compact libraries have been introduced to reduce screening costs and accommodate more advanced screening strategies as well as models with limited growth, such as primary cells and patient derived organoids: Gattinara reflects the same design approach as Brunello to select two guides per gene ^24^, while MinLibCas9 features a guide-per-gene count (quota) of two as well, prioritizing guides featured in Project Score to ensure backward compatibility and integration with Project Score’s existing screens ^25^.

Several considerations motivate developing a new Cas9 CRISPRko genome-wide library. First, updates to gene annotations over time cause deprecation of existing libraries; in some cases, gene-targeting guides designed against prior annotations map to sites that fall outside of protein coding regions, while novel genes discovered in recent annotations are not targeted by previous libraries ^26^. Additionally, the development of the Rule Set 3 model of on-target activity, which captures the interactive effect of an expansive set of guide and target site features as well as the choice of tracrRNA, provides the opportunity to design more effective libraries ^27^. Likewise, recent approaches to avoid guides that target genomic sites with high SNP frequencies can be incorporated in library design to minimize false negative rates across cell models with various ancestries ^28^.

Libraries can further improve efficiency through a strategic balance of on-target and off-target considerations. To date, numerous *in silico* methods have sought to reliably predict SpCas9 guide efficacy at the target site ^15,23,27,29–31^ as well as the probability of activity at putative off-target sites ^15,32–36^. However, much less attention has been paid to the balance of these criteria for the ultimate selection of guides to include in a library ^16^. While over-prioritizing a guide’s propensity for on-target activity may result in the inclusion of promiscuous guides, avoiding those with only a slight chance of off-target activity may eliminate the guides that effectively perturb the target gene. Here we present a data-driven approach to detect guides with sufficient promiscuity to justify exclusion from a Cas9 CRISPRko screen, thus maintaining the representation of guides with strong on-target activity and tolerable levels of off-target activity. We employ this approach for the design of an optimized genome-wide SpCas9 CRISPRko library, Jacquere.

## RESULTS

### Establishing patterns of Cas9 off-target activity from GUIDE-seq data

Experimental methods to profile guide activity across the genome have enabled precise quantification of off-target activity for individual guides, yet such assays are not feasible to implement for the evaluation of all candidate guides for a CRISPR screen ^37,38^. As a complement to these efforts, we previously screened guides mismatched to CD33 to generate a cutting frequency determination (CFD) matrix, which reports the rate at which guides remain active despite a mismatch of each possible type and position along the target DNA ^15^. The CFD matrix demonstrates that mismatches to the target in guide positions 4-20 and deviations from the canonical PAM generally disrupt guide activity, whereas mismatches in the PAM-distal positions 1 through 3 are often tolerated. We thus define the former window as the specificity-defining region (SDR) (Figure 1a).

**Figure 1:**
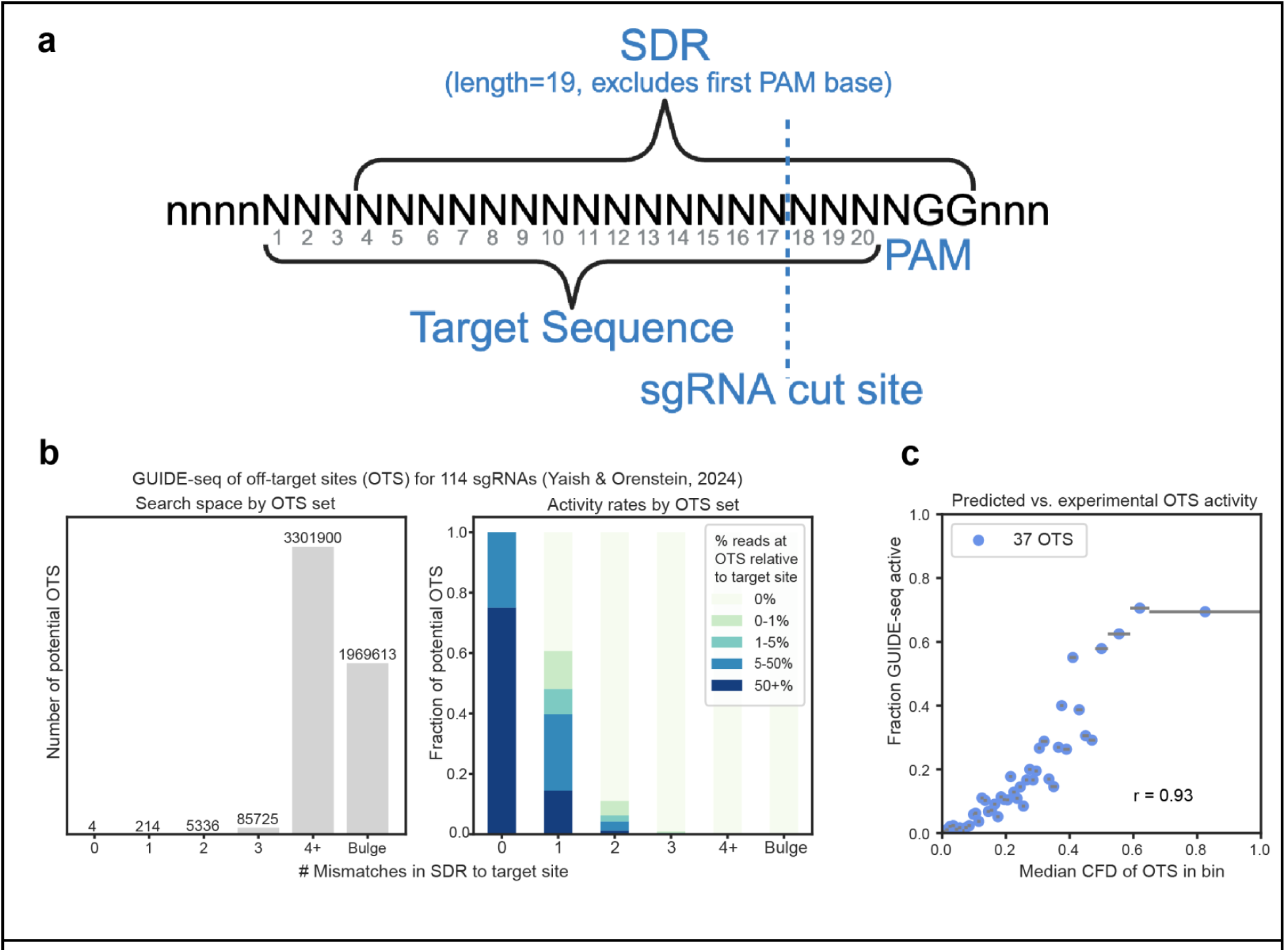
Benchmarking predictions of Cas9 CRISPRko guide behavior at putative off-target sites with GUIDE-seq data. a) The Specificity Defining Region (SDR) is the window we employ in this manuscript and in the CRISPick library design tool to determine the number of mismatches between a guide and a putative off-target site. Mismatches at positions 4 through 20 along the guide and to the canonical PAM are counted towards this total. b) Patterns of Cas9 nuclease activity at off-target sites as revealed by GUIDE-seq data. Left: Off-target site counts indicate the total number of potential off-target sites across the 114 guides in the dataset, stratified by the maximum number of mismatches in the SDR tolerated in the off-target search. “Bulge” off-target sites represent those with 4+ mismatches in the SDR, yet the ability to reduce this mismatch count with a single nucleotide shift within the sequence. Right: the fraction of each set of off-target sites as reported on the left in each range of activity rates, which are measured by the percent of GUIDE-seq reads at the off-target site relative to the guide’s on-target site. c) Association between CFD scores and activity at all predicted off-target sites for the 114 guides in the GUIDE-seq dataset. Off-target sites that exceed two mismatches in the SDR are excluded due to their low propensity for activity. Off-target sites are binned into groups of 37 by their CFD score, and those with at least 1% as many GUIDE-seq reads as the on-target site are classified as active. For each bin, the median CFD value indicates the predicted activity rate, and the fraction of active off-target sites indicates the experimental activity rate, yielding a Pearson correlation of 0.93. Horizontal error bars capture the minimum and maximum CFD score within each bin.

The CFD score, namely the probability (scaled 0 - 1.0) that a guide will remain active despite mismatches at a putative off-target site, is the product of tolerance rates from the CFD matrix corresponding with each mismatch. For example, an off-target site with several PAM-proximal mismatches to the guide will have a CFD score near 0 to indicate low threat of off-target activity. Despite its remarkable simplicity and ease of implementation, CFD score has been found to outperform hypothesis-driven methods and approaches the precision of state-of-the-art deep learning models in its predictions of off-target activity against experimentally validated guides ^39^.

To refine predictions of Cas9 promiscuity, we refer to GUIDE-seq, an assay that tags, amplifies, and sequences sites of double-stranded breaks with the use of an indicated guide, thereby yielding read counts proportional to the rate of guide activity at an off-target site ^37^. We leverage a recent work that aggregates a large corpus of GUIDE-seq experiments for 114 unique guides across primary CD4^+^/CD8^+^ T cells, U2OS, and HEK293 cell lines^40^. Overall, we observe that numerous mismatches in the SDR tend to inhibit Cas9 activity; at putative off target sites with 0, 1, 2 or 3 mismatches, guides are active – defined as having at least 1% as many GUIDE-seq reads as the on-target site – at a rate of 100%, 43%, 5.7%, and 0.4%, respectively (Figure 1b). Additionally, we find that numerous-mismatch off-target sites for which a shift in alignment by one position would resolve mismatches show no increase in activity, suggesting that off-target activity as a result of single nucleotide bulges in the DNA or RNA are unlikely (Figure 1b).

These results suggest that, since the vast majority of putative off-target sites with three or more mismatches to the guide in the SDR are false positives, it is appropriate to disregard such sites when evaluating guides for genome-wide screens. In addition to the fact that guide promiscuity is less detrimental in this context relative to therapeutic applications of CRISPR technology, omitting effective guides on the false pretense of off-target activity diminishes the informative potential of such screens. Moreover, the computational complexity and memory usage implicated in the search for these vastly inactive off-target sites is not merited; while searching for all off-target sites with up to two mismatches to a single guide involves 1,594 queries against the genome, this figure increases to 27,760 queries when extending to three mismatches and 341,716 queries when extending to four. Repeated across the 5,273,116 candidate Cas9 CRISPRko guides in total among protein-coding genes, reducing the off-target search even from 3 to 2 mismatches would not only reduce the false positives involved in guide evaluation but also improve the efficiency of library design by eliminating nearly 138 billion queries against the genome (see Methods).

Lastly, beyond revealing the scope of potential Cas9 CRISPRko off-target sites, this large-scale GUIDE-seq dataset informs the extent to which the CFD score reflects experimentally determined rates of off-target activity. Upon calculating the CFD score between the 114 guides and all off-target sites with up to two mismatches, grouping off-target sites by CFD score, and identifying the fraction of each group that are active, we find that CFD score correlates with rates of GUIDE-seq activity (Pearson R=0.93)(Figure 1c). We therefore deem it appropriate to leverage the CFD score in off-target avoidance strategies for Cas9 CRISPRko library design.

### Calibrating a classifier to detect guide promiscuity

Promiscuous guides primarily confound CRISPR knockout screens by introducing sufficient double-stranded breaks across genomic sites to activate DNA damage response, thereby reducing cell fitness in a target-independent manner ^32,41,42^. Notably, numerous double-stranded breaks contribute to the antiproliferative effects of a guide in an additive manner ^43^. The specificity score developed for the GuideScan tool serves as a breakthrough in the efforts to quantify the cumulative off-target activity of a guide; namely, it is the reciprocal of the sum (+1 to avoid division by zero) of CFD scores for all off-target sites with up to 3 mismatches in the 20 nt guide sequence ^44^. This metric has been found to outperform alternatives, such as the MIT score, at distinguishing guides with extensive off-target activity ^45^. The manuscript accompanying the release of GuideScan2 advises that libraries omit guides exceeding a specificity score of 0.2 ^16^. However, the set of off-target sites included in the specificity score is expansive in light of evidence that Cas9 has minimal activity at numerous-mismatch off-target sites (Figure 1b), so here we attempt to develop an optimized classifier for promiscuous guides.

To train our classifier, we leverage the Rule Set 3 Validation set viability screens ^27^, which tile guides across nonessential genes and identify promiscuous guides as those that deplete in the population. We preserve the framework of representing each guide as the sum of CFD scores at potential off-target sites and vary the maximum number of mismatches to the guide in the SDR in the search for off-target sites (including those with a noncanonical PAM) to assess its impact upon predictions of guide promiscuity (Figure 2a). The use of the SDR to determine mismatch count improves the chance of incorporating active off-target sites while avoiding excessive computational time and complexity. For example, to identify all off-target sites with up to one mismatch in the SDR (19nt total) results in 58 queries against the genome. When incorporating the first three guide positions in the mismatch count, one would need to search for sites with up to 4 mismatches to guarantee the detection of high-risk off-target sites, namely those that only have a single mismatch in the SDR, which results in 592,515 queries against the genome.

**Figure 2:**
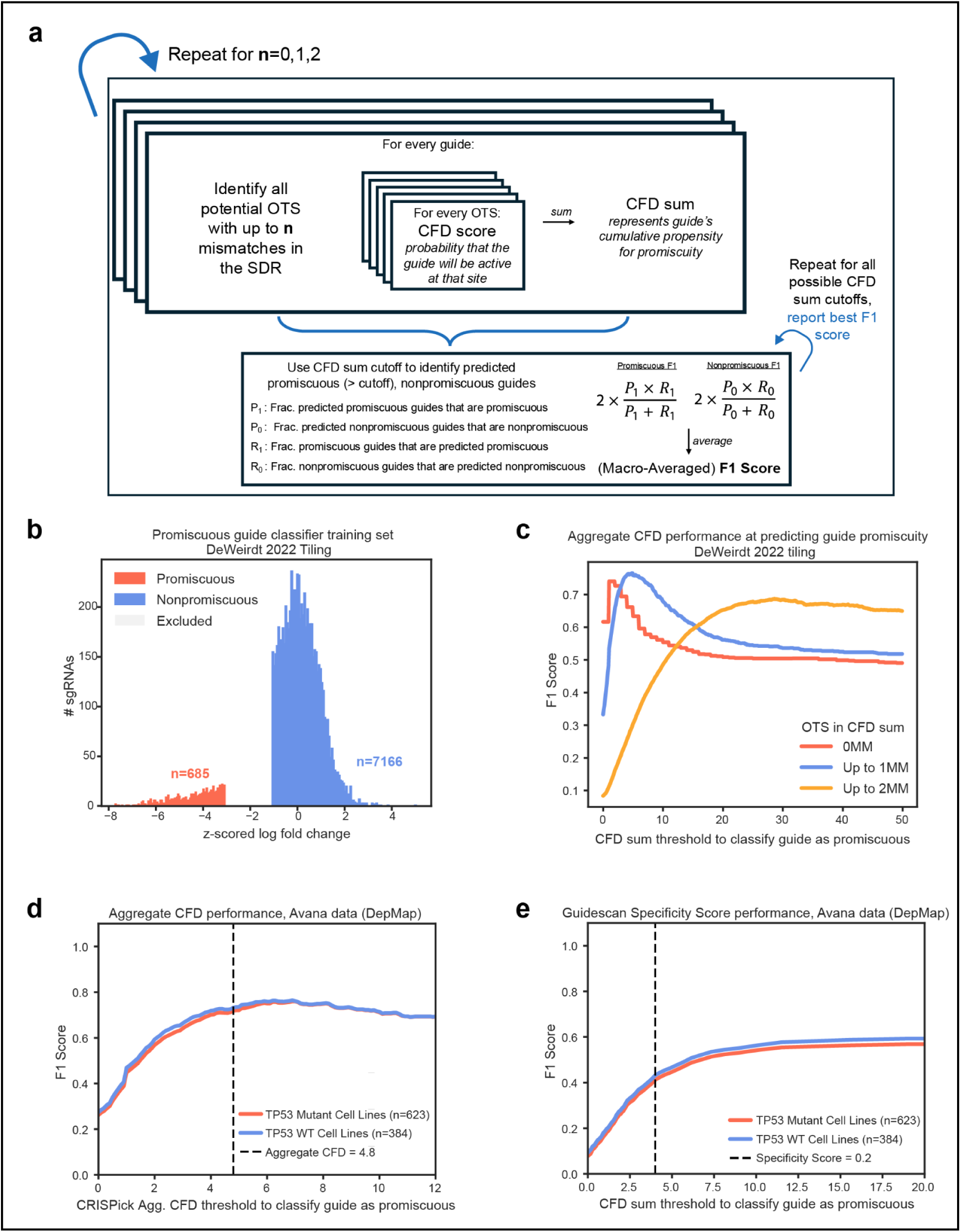
The development of a classifier to detect guides that introduce an antiproliferative number of double-stranded breaks at off-target sites. a) The protocol that is implemented to develop a classifier for predicting guide promiscuity. The off-target potential of each guide in the training set is quantified by adding the CFD scores at each putatitve off-target site. The Macro-Averaged F1 score is then used to identify the CFD sum that best distinguishes true promiscuous and non-promiscuous guides as determined from screening data. This process is repeated, varying the search space of off-target sites to those with up to 0, 1 or 2 mismatches to the guide in the SDR, to identify the scope of off-target sites that yields the highest Macro-Averaged F1 score and thus recognizes true promiscuous guides most effectively. b) The distribution of nonessential-targeting guides in the DeWeirdt 2022 tiling set. For the training of a classifier of guide promiscuity, promiscuous guides are identified as those with a z-score below −3 relative to the intergenic controls, and non-promiscuous guides have a z-score above −1. c) The training performance of CRISPick Aggregate CFD as a function of the threshold employed to classify a guide as promiscuous. Performance is stratified by the off-target sites included in the sum. d) The validation performance of CRISPick Aggregate CFD as a function of the threshold employed to classify a guide as promiscuous. Performance is stratified by cell line TP53 status. All off-target sites with up to one mismatch in the SDR are included in the CFD sum. The black dotted line indicates the threshold established from Aggregate CFD training to classify guides as promiscuous. e) The validation performance of the GuideScan specificity score at predicting guide promiscuity. The x-axis reports the reciprocal of the specificity score (with 1 subtracted) to indicate the CFD sum of off-target sites for ease of comparison with CRISPick Aggregate CFD. The black dotted line corresponds with a specificity score of 0.2 to demonstrate the threshold suggested by the authors of GuideScan2 to classify guides as promiscuous in library design. The maximum possible F1 scores of 0.59 for TP53WT and 0.57 for TP53 Mutant cell line data exceed the range of the plot.

To tune the classifier, we assess model performance with the Macro-Averaged F1 score (hereinafter abbreviated as F1 score), which identifies a random binary classifier as F1= 0.5 and a perfect classifier as F1=1.0 (Figure 2a). Unlike the standard F1 score, the macro-averaged F1 score computes the F1 score separately with promiscuous and non-promiscuous guides as the positive class and reports the average, thereby evenly prioritizing accurate classification of both groups ^46^. The use of this metric avoids bias introduced by the underrepresentation of promiscuous guides in the training data (Figure 2b); alternative metrics such as the area under a receiver operating curve (ROC-AUC) can yield misleading estimates of performance for data with imbalanced class sizes ^47^.

We find that the CFD sum of off-target sites with up to 0, 1, or 2 mismatches in the SDR optimally predicts promiscuous guides at F1 scores of 0.74, 0.77, and 0.69, respectively, at CFD sum thresholds of 1.0, 4.8, and 28.6 (Figure 2c). The CFD sum therefore achieves the most effective predictions of guide promiscuity when only considering off-target sites with up to one mismatch, a result that is consistent with the observation from GUIDE-seq data that off-target sites with numerous mismatches are largely false positives (continuing to use the 1% cutoff to define an off-target site as active, as described above). We refer to the CFD sum with these specifications as the CRISPick Aggregate CFD, building off of the term “Aggregate CFD” introduced in prior works ^10,45^.

To validate the performance of CRISPick Aggregate CFD, we apply this classifier to screening data of nonessential-targeting guides in the Avana library, which features simplified avoidance of off-target activity in its design, prior to the development of the CFD score ^15^. We stratify the evaluation of this metric for TP53 mutant (n=623) and wildtype (n=384) cell lines to assess if the increased sensitivity to double stranded breaks of the latter impacts performance ^48^. At its optimized classifier threshold of 4.8 from the training set, CRISPick Aggregate CFD has an F1 score of 0.73 and 0.72 for TP53 wildtype and mutant cell lines respectively (Figure 2d).

Importantly, the stable performance of CRISPick Aggregate CFD regardless of cell line TP53 status suggests that this model is widely applicable for the evaluation of Cas9 CRISPRko guides.

While CRISPick Aggregate CFD and the GuideScan specificity score both leverage CFD scores to compute a weighted sum across off-target sites, the latter includes off-target sites with up to 3 mismatches in the 20nt guide sequence whereas CRISPick Aggregate CFD only considers those with up to 1 mismatch in the SDR^16^. Consequently, the GuideScan specificity score incorporates more inactive off-target sites; in the aforementioned GUIDE-seq data, 87.9% of off-target sites flagged by the GuideScan specificity score are inactive in contrast to the 56.3% of off-target sites identified by CRISPick Aggregate CFD (Supplementary Figure 1). These inactive off-target sites inflate the GuideScan specificity score and compromise its performance at distinguishing promiscuous guides, yielding optimized F1 scores of 0.59 for TP53 Wildtype and 0.57 for TP53 Mutant cell line data. The F1 score further drops to 0.43 and 0.41 when using the suggested specificity score threshold for library design of 0.2 (Figure 2e). Despite high detection rates of promiscuous guides (97.9% for TP53 WT, 92.7% for mutant), this cutoff yields a suboptimal rate of identifying true non-promiscuous guides (41.9% for TP53 WT, 42.4% for mutant), which is in part driven by its automatic promiscuous classification of all guides with perfect or multiple single-mismatch off-target sites (considering all positions along the 20nt guide). We therefore recommend the use of CRISPick Aggregate CFD for recognizing guides that introduce an antiproliferative number of double stranded breaks at off-target sites.

### Design of Jacquere, a Cas9 CRISPRko genome-wide library with improved guide efficiency

#### An updated landscape of target genes

The RefSeq and GENCODE databases, which are curated by NCBI and Ensembl, respectively, both provide reliable estimates of the set of human protein coding transcripts. We further recognize the validity of the CHESS 3 database, which has determined the set of human genes and their transcripts through robust processing of RNA sequencing data from the GTEx project ^49^. Since novel genes identified in previous iterations of CHESS were subsequently added to GENCODE, RefSeq, and the MANESelect consensus set (with each database annotating 53, 23, and 5 novel CHESS genes respectively), CHESS 3 may be indicative of genes to be added to the more widely-recognized databases in the future. To reflect the developing understanding of the human protein-coding transcriptome, the Jacquere library targets the comprehensive set of genes identified by the RefSeq, GENCODE, and CHESS databases, referring to the primary transcript in all cases (Figure 3a). While Jacquere targets a contemporary set of protein coding genes at present, web tools such as CRISPick will continue to be an up-to-date source of information as genome annotations continue to evolve.

**Figure 3:**
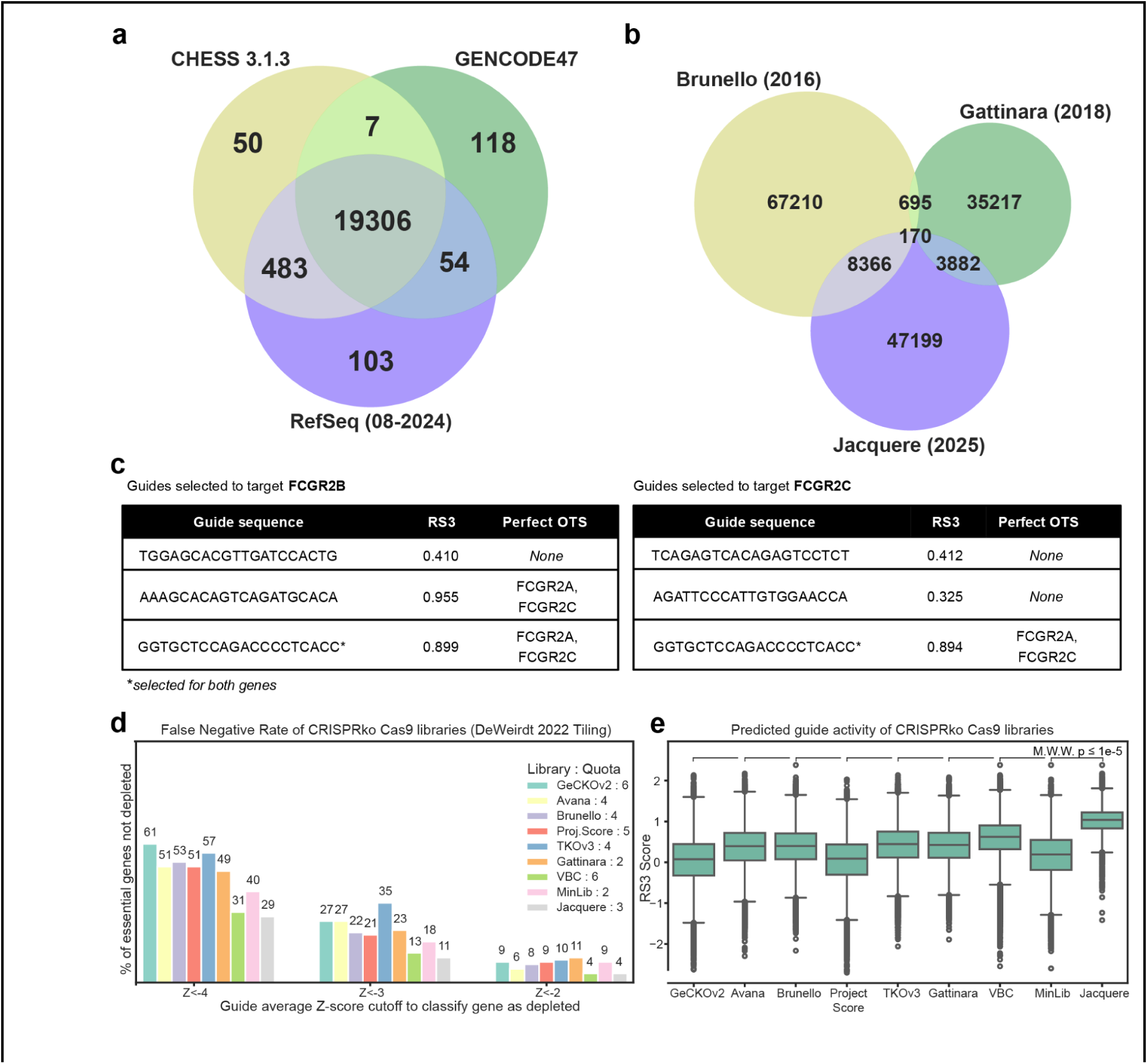
Composition of Jacquere and comparison across Cas9 CRISPRko Genome-wide Libraries. a) The overlap of genes annotated by RefSeq 08-2024, GENCODE 47, and CHESS 3.1.3. Genes may overlap between databases if they are annotated by distinct symbols or IDs by their respective databases but correspond to the same HGNC symbol. Circles representing each subset and their degree of overlap are not to scale. b) The overlap between gene-targeting guide sequences present in comparable libraries released by the Genetic Perturbation Platform: Brunello (2016), Gattinara (2018), and Jacquere (2025). Circle sizes are scaled to represent the size of each library and degree of overlap. c) A summary of the guides targeting FCGR2B and FCGR2C in Jacquere to exemplify the effect of the guide selection strategy on the design of paralogous genes. The guide selected for both genes (denoted in the figure with the * symbol) yields two distinct RS3 scores since this model incorporates target-specific features. d) An assessment of false negative rates across various genome-wide Cas9 CRISPRko libraries as measured by the percent of essential-targeting genes that fail to yield substantial average guide-level depletion. This analysis is conducted with the DeWeirdt 2022 tiling library screening data, and z-scores are calculated relative to the mean and standard deviation of intergenic controls ^27^. e) The distribution of RS3 scores representing the predicted on-target activity of guides across Cas9 CRISPRko genome-wide libraries. Boxes depict 25th (Q1) and 75th (Q3) percentiles as minima and maxima and the center represents the median; whiskers identify outlier points by depicting Q1 - 1.5*IQR and Q3 + 1.5IQR, where IQR represents the range between Q1 and Q3.

#### Compact library size enables broad application and reflects advances in guide selection

Genetic screens employ numerous guides per gene to improve the chance of successful perturbation and recognize off-target effects through the presence of outlier guides. When previously designing Brunello, we identified that four guides targeting each gene are required to retain a satisfactory signal-to-noise ratio ^24^. The design of Jacquere reflects a revised strategy to prioritize guide efficacy, as evidenced by distinct guide selections in Jacquere from libraries designed with the prior approach (Figure 3b), thereby enabling the use of fewer guides to infer gene function. For this reason, the Jacquere library features a quota of 3 guides per gene, a choice that we discuss in greater detail below.

#### Balanced prioritization of on-target and off-target guide properties

Efficacy at the on-target site is the first priority when selecting guides for inclusion in Jacquere. Upon excluding guides that explicitly violate specifications discussed below, guides are selected to target each gene in order of their potential for on-target activity as determined by the RS3 score. In addition, we preferentially select guides with a RS3 score of 0.2 or above, which is associated with depletion of essential-targeting guides (FDR <0.05) in Brunello screening data (χ^2^(1,N=5061)=32.8, p=<0.001) ^50^. This cutoff is the last criteria that can be relaxed in guide selection if necessary to meet the quota for a gene, thereby ensuring that guide efficacy is not compromised in light of additional design considerations (Supplementary Data 1).

For off-target avoidance, the design of Jacquere prohibits the inclusion of guides that exceed a CRISPick Aggregate CFD score of 4.8 since they are expected to generate an antiproliferative number of double stranded breaks at off-target sites. A second, lower probability mechanism of deleterious guide behavior involves activity at a single off-target site with a phenotypic consequence that can be detected by the screening readout ^32^. To avoid this source of error in a context-agnostic manner, we preferentially select guides without any CFD=1.0 off-target sites, which indicates a predicted 100% chance of off-target activity as determined by the 17 nucleotides preceding the PAM sequence. CFD=1.0 off-target sites are tolerated if necessary to meet a gene’s quota, avoiding such sites in protein coding regions when possible. This flexible approach enables balance between off-target avoidance and guide efficacy while still achieving consistent gene coverage.

#### Additional guide selection criteria

A recent work has estimated that 1.2-2.5% of guides in genome-scale Cas9 CRISPRko libraries have reduced or eliminated cutting efficiency as a result of germline variants at the target site, a pattern notably present in cell lines sourced from individuals with African ancestry ^28^. To mitigate the risk of cell model SNPs impeding guide activity, the design of Jacquere avoids guides that target sites with high variability in the human population, as defined by a gnomAD variant frequency of above 5%. Since variant frequencies from the total population may not capture variants common to the African/African American subgroup, which is disproportionately underrepresented in the human reference genome, library design additionally avoids target sites with a variant frequency of at least 12.5% in this group^51^. As a result, only 0.08% of targeting guides in Jacquere exceed either of these thresholds.

More generally, guide selection for Jacquere leverages existing criteria in CRISPick for practical library design. Among all candidate guides for each gene, we preferentially select those that target between 5-65% along the protein coding region of the target gene to improve the chance that double-stranded breaks disrupt protein function. Lastly, while CRISPick targets the canonical transcript for each gene, we preferentially select guides spaced at least 5% of the length of the protein coding region apart (in amino acid space) to improve the chance that guides target various exons and thus isoforms.

#### Effect of library design approach on paralogs

Paralogous genes are highly relevant for functional association studies, by one estimate comprising 80% of disease associated genes ^52^. It is therefore critical to ensure strategic guide selection for such genes. To accommodate future updates to genome annotations that will inevitably modify the comprehensive set of paralog families, we apply the same library design approach for all genes and ensure that this strategy yields optimal guide selections for paralogs and nonparalogs alike.

Existing studies discourage targeting shared sequences across paralogs, cautioning that the use of multi-target guides may cause hits to arise in a cell-line dependent manner due to the antiproliferative effects of numerous double stranded breaks ^43^. However, avoiding multimapping guides may inhibit the selection of effective guides for paralogous genes; in addition to restricting the set of candidate guides for paralogs whose protein-coding sequences are largely identical, this design choice disproportionately eliminates guides that target conserved regions, which are more likely to be effective ^27^. Indeed, there are 484 paralogous genes that are targetable only with multitarget guides (Ensembl v113), which extends to 603 genes when restricted to effective guides (RS3 >0.2). Tolerating multimapping guides thus promotes the coverage of paralogous genes with guides with strong on-target activity, thereby justifying the decision to not exercise absolute exclusion of guides with perfect off-target sites. The risk of antiproliferative effects introduced by this design choice is mitigated through the exclusion of guides that violate the CRISPick Aggregate CFD threshold of 4.8.

It is additionally important that the design strategy for Jacquere allows a single guide to be selected for more than one target gene. As an example, the paralogous Fc gamma receptors FCGR2B and FCGR2C feature 82.5% sequence similarity. Following the design approach applied to all targets in Jacquere, the guides first selected for each gene feature RS3 scores of at least 0.2, target sites 5-65% along the protein, and do not have CFD=1.0 sites in the coding region of other genes. This step incorporates reasonably effective guides that target only one of the paralogs, allowing the library to identify potential discrepancies in gene function. To meet the quota for these genes, the criteria are relaxed to tolerate CFD=1.0 off-target sites in protein coding regions, and the candidate guides with the highest RS3 scores that target 5-65% along the protein are selected. The best candidate guide at this step is the same for FCGR2B and FCGR2C, resulting in these two genes sharing a guide (Figure 3c). Since candidate guides that effectively target these genes uniquely have been exhausted, any remaining candidate guides for these genes would target both, just with a lower propensity for on-target activity. Thus, to avoid the unnecessary inclusion of ineffective guides when each of these genes is targeted at quota level, assigning this guide to target both genes is appropriate.

#### Comparison to existing Cas9 CRISPRko genome-wide libraries

We have previously demonstrated that the use of four guides to target each gene optimizes screening performance with existing library design approaches ^24^. However, improvements to library design pose an opportunity to achieve satisfactory screening performance at a reduced quota. With the use of three guides per gene, the guide selections resulting from the Jacquere library design approach feature the lowest false negative rate as shown with screening data for 201 essential genes (Methods) (Figure 3d). The VBC library yields the closest false negative rates to Jacquere (albeit with the use of six instead of three guides per gene), reflecting the merit of employing guides favored to introduce loss-of-function phenotypes based on the amino acid composition of the expected edit. We anticipate the performance of libraries designed by leveraging VBC scores could improve with the use of Rule Set 3 instead of Rule Set 2 to model on-target activity ^23^.

The official Jacquere library targets 20,550 genes, of which 99.5% meet the quota of three guides per gene (Supplementary Figure 2a). The library contains 60,617 unique guides, including 900 intergenic and 100 non-targeting guides that feature comparable predicted on-target activity (median RS3 1.11 compared to 1.04 for gene-targeting guides) and off-target risk (Supplementary Figure 2b). 95.0% of guides meet the top standards of design criteria (Supplementary Data 1), and only 2.0% of guides map to more than one protein coding gene (with respect to Ensembl v113 annotations). Designs are supplied to indicate each guide sequence in Jacquere, the gene(s) for which the guide is selected to target, and the full set of genes to which the guide maps (Supplementary Data 2). We additionally provide designs that include a fourth guide selection per target for relevant screening applications (Supplementary Data 3).

Using GENCODE47 as reference, Jacquere achieves higher coverage of the genome in comparison to existing Cas9 CRISPRko genome-wide libraries (Table 1) ^26^. Jacquere additionally demonstrates a higher distribution of RS3 scores (Mann Whitney one-sided U-test. p<0.00001) and lower representation of guides with high CRISPick Aggregate CFD or CFD=1.0 off-target sites, demonstrating its ability to prioritize guide efficiency without compromising off-target avoidance (Figure 3e, Table 1). Importantly, while the average CRISPick Aggregate CFD of Jacquere guides is not the lowest among libraries, Jacquere is the only library with 0 guides exceeding a score of 4.8, demonstrating the strategic design choice to only prioritize off-target avoidance for guides with sufficiently high risk of off-target activity to reduce cell viability.

**Table 1:** Information for each genome-wide Cas9 CRISPRko screening library compared in this manuscript. The genomic coverage indicates the percent of GENCODE47 genes that are targeted by 0 guides. The CRISPick Aggregate CFD composition and percentage of guides with 100% CFD probability off-target sites (OTS) inform the off-target propensity of guides in each library.

| Library | Release Year | #Guides per gene | PMID | Addgene ID | % of GENCODE47 genes untargeted | Aggregate CFD |  | CFD 1.0 OTS |  |
| --- | --- | --- | --- | --- | --- | --- | --- | --- | --- |
|  |  |  |  |  |  | Mean | % >4.8 | % in full genome | % in protein coding regions |
| GeCKOv2 | 2014 | 6 | 25075903 | 1000000048 | 2.72 | 3.11 | 2.9 | 14.3 | 3.4 |
| Avana | 2016 | 4 | 26780180 | NA | 3.48 | 2.13 | 4.6 | 15.4 | 3.3 |
| Brunello | 2016 | 4 | 26780180 | 73178 | 2.23 | 0.79 | 2.2 | 6.2 | 3.3 |
| Proj.Score | 2016 | 5 | 27760321 | 67989 | 6.98 | 0.64 | 1.4 | 9.8 | 2.4 |
| TKOv3 | 2017 | 4 | 30970261 | 90294 | 7.57 | 0.76 | 1.1 | 11.1 | 1.3 |
| Gattinara | 2018 | 2 | 32029722 | 136986 | 1.33 | 1.01 | 3.1 | 13.1 | 3.8 |
| VBC | 2020 | 6 | 32514112 | NA | 1.73 | 1.13 | 2.8 | 12.6 | 1.8 |
| MinLib | 2021 | 2 | 33478580 | 164896 | 3.13 | 0.58 | 1.2 | 6.3 | 2.1 |
| Jacquere | 2025 | 3 | NA | tbd | 0.61 | 0.78 | 0 | 3.2 | 2.2 |

### Jacquere screening performance

To validate the performance of Jacquere, we conducted viability (depletion) screens in the A549 (lung) and A375 (melanoma) cancer cell lines, the latter of which we previously screened with the Brunello library ^50^. With the use of a vector that delivers both Cas9 and the guide RNA (“all-in-one”), we observe consistent performance across biological replicates (Pearson correlation coefficient = 0.93, 0.95 for A375, A549 respectively) (Figure 4a). Jacquere offers improved distinction between guides targeting essential and nonessential genes over the Brunello Library, as demonstrated by a receiver operating characteristic area under the curve (ROC-AUC) of 0.96 and 0.92 for Jacquere and Brunello respectively (Figure 4b). Further, a precision-recall curve reveals the simultaneously precise and sensitive detection of essential genes by the Jacquere library (AUC=0.98, 0.95 for Jacquere and Brunello, respectively, screened in A375) (Figure 4c). The Jacquere library maintains strong performance when the guide and Cas9 are delivered to the cells separately (Supplementary Figure 3).

**Figure 4:**
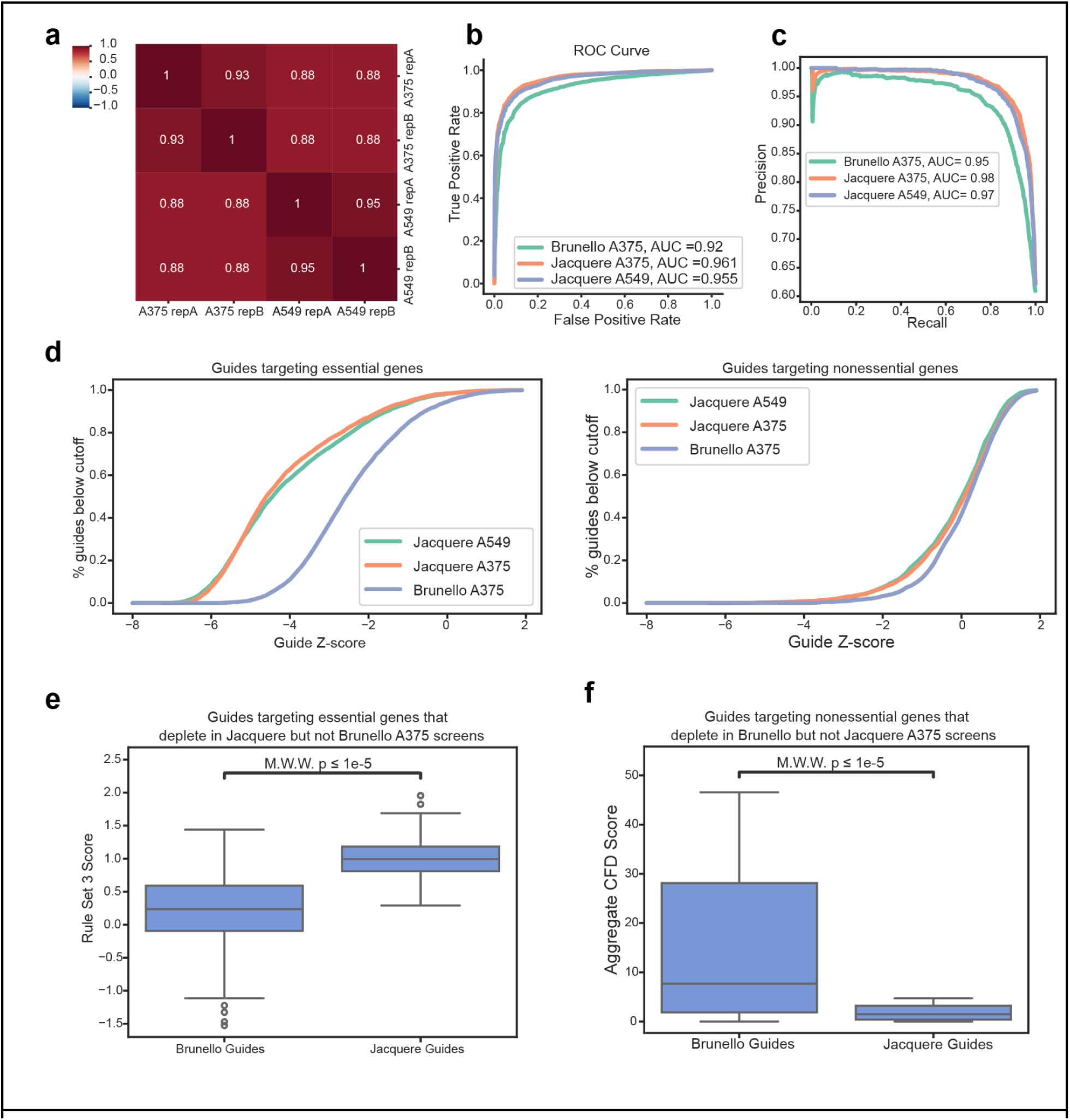
Assessment of the Jacquere library screening performance. a) The Pearson correlation between biological replicates and across cell lines for the Jacquere Library screened with the pRDA_734 vector. Correlation is calculated upon the log-fold change in guide abundance at Day 21 post-transduction relative to pDNA abundance. b) Receiver operating characteristic analysis of cell viability screening data for the Jacquere library (pRDA_734 vector) and Brunello Library screened in A375 cells. False positive rates are determined by non-essential genes and plotted against the true positive rate, which is determined by essential genes. c) Precision recall curve to compare the classification of essential genes between Jacquere and Brunello screening data. d) Stratified analysis of false positive rates and false negative rates in screening data of the Jacquere and Brunello libraries. Guide z-scores are calculated relative to the mean and standard deviation of intergenic control guides, and the plots demonstrate the percentage of guides targeting essential (left) or nonessential (right) genes that deplete at the z-score cutoff as indicated by the x-axis. e) Rule Set 3 sequence+target (Chen tracrRNA) on-target efficacy scores for guides selected by each library to target the essential genes that deplete in Jacquere but not Brunello screening data, thus representing false negatives unique to the Brunello library. Boxes depict 25th (Q1) and 75th (Q3) percentiles as minima and maxima and the center represents the median; whiskers identify outlier points by depicting Q1 - 1.5*IQR and Q3 + 1.5IQR, where IQR represents the range between Q1 and Q3. f) Aggregate CFD Scores for guides selected by each library to target the nonessential genes that deplete in the Brunello but not Jacquere screening data, thus representing false positives unique to the Brunello library. Boxes depict 25th (Q1) and 75th (Q3) percentiles as minima and maxima and the center represents the median; whiskers identify outlier points by depicting Q1 - 1.5*IQR and Q3 + 1.5IQR, where IQR represents the range between Q1 and Q3.

Whereas both the Jacquere and Brunello libraries feature comparable false positive rates, as assessed by depletion of guides targeting nonessential genes, the Jacquere library features a markedly reduced false negative rate, with improved depletion of essential targeting guides (Figure 4d). Upon aggregation of guides targeting the same gene, 130 common essential genes fail to deplete (at FDR<0.05) when screened with Brunello, and 97 of these essential genes are recovered as a hit when screened with Jacquere. These recovered hits are associated with the prioritization of RS3 scores in the design of the Jacquere library; the Jacquere guides that target these genes indeed feature higher RS3 scores than those used in the Brunello library (Mann Whitney two-sided U-test. p<0.001, Figure 4e). The screening results of these two libraries additionally imply the merit of avoiding guides with Aggregate CFD scores; among nonessential genes that deplete (FDR<0.05) in the Brunello but not the Jacquere library, the guides targeting these false positives of the Brunello library have lower Aggregate CFD scores in the Jacquere library (Mann Whitney two-sided U-test. p<0.001, Figure 4f). Overall, this comparison shows that the guide selection principles we describe in this manuscript result in a library with excellent performance.

### Hit identification for genetic screens

Factors contributing to false positive and false negative rates of CRISPR screens extend beyond what can be controlled in library design. Certain genes may lack PAM-adjacent sequences that are specific and have a high propensity for Cas9 guide activity, such as the aforementioned 603 genes that are only targetable by ineffective (RS3<0.2) or multimapping guides, resulting in the necessary inclusion of suboptimal guides in the library. Further, guides favored for on-target activity may still fail to produce a loss of function allele due to SNPs in the cell model or the introduction of an innocuous in-frame mutation ^23^. Thus, even the most meticulous library design does not eliminate the need for strategic downstream analysis to determine gene-level effects from guide-level evidence.

For screening readouts that profile each guide with a single numeric value, e.g. guide depletion for viability screens, there exist several methods to aggregate guides with the same target gene. While there exists a plethora of methods that incorporate advanced features, such as prior assumptions^53^ or the ability to correct for copy number variation in the cell line employed ^54^, here we focus upon methods that require no input aside from the raw screen results and genes targeted by each guide. The Z-score method, as exemplified in the analysis of the Jacquere library screen, entails representing each guide as the number of standard deviations from the mean with respect to negative control guides and aggregating the scores per gene with Stouffer’s method. Notably, this approach is conceptually straightforward, convenient to implement in a screen analysis pipeline, and demands a negligible amount of computing time.

We additionally examine the hit-calling methods MAGeCK RRA and MAGeCK MLE ^55,56^. Much like the bulk RNA sequencing differential expression methods edgeR and DESeq2, the MAGeCK methods model guide effects with a negative binomial distribution to account for over-dispersion, namely, the tendency of read counts generated from high-throughput sequencing data tend to yield higher variance at higher counts ^57,58^. MAGeCK RRA (Robust Rank Algorithm) evaluates genes with the assumption that, upon ranking guides by their change in count relative to the initial quantity, genes with no selective pressure will have an even distribution of targeting guides across the rank. As an effort to prevent ineffective guides from introducing false negatives, the algorithm allows the guide with the most severe rank to determine if a gene is a hit. In contrast, MAGeCK MLE (Maximum Likelihood Estimation) determines a gene’s effect from its magnitude of selective pressure, ascribing greater weight in this estimate to guides postulated to be effective using an expectation maximization procedure. While MAGeCK MLE is more recent, MAGeCK RRA runs faster, so both methods are still in widespread use to identify hits from CRISPR screens.

To provide guidance on navigating the choice of a downstream analysis method, here we apply the Z-Score method, MAGeCK RRA, and MAGeCK MLE to Jacquere and Brunello screening data. Both MAGeCK hit-calling methods involve a statistical test to determine the significance of the observed response to the gene’s loss of function. However, previous works have found such p-values to be miscalibrated, subjecting screens to high false discovery rates ^59^. We therefore calculate empirical p-values from MAGeCK MLE beta scores and MAGeCK RRA enrichment scores with pseudogenes constructed from intergenic controls as the null distribution (see Methods, Analysis of Jacquere and Brunello screening data). Upon Benjamini-Hochberg correction for multiple testing, we recognize the hits identified from each method as those with a false discovery rate (FDR) of below 5%.

The FDRs resulting from all three methods are able to effectively distinguish known essential from nonessential genes, with improved performance upon Jacquere screening data (Figure 5a). An FDR cutoff of 0.05 demonstrates effective control of the fraction of essential genes that each method fails to detect as depleted as well as the fraction of nonessential genes falsely identified as depleted, suggesting that this is an appropriate threshold to determine hits in this context (Figure 5b). The set of genes that emerge as hits from a screen are somewhat subject to library composition, as exemplified by the slight distinction in the hits identified by analysis of the Jacquere and Brunello libraries, and this effect is consistent regardless of the hit-calling method employed (Figure 5c). Further, all three methods mostly identify the same set of essential genes as significantly depleted within both the Jacquere and Brunello screen (Figure 5d,e).

**Figure 5:**
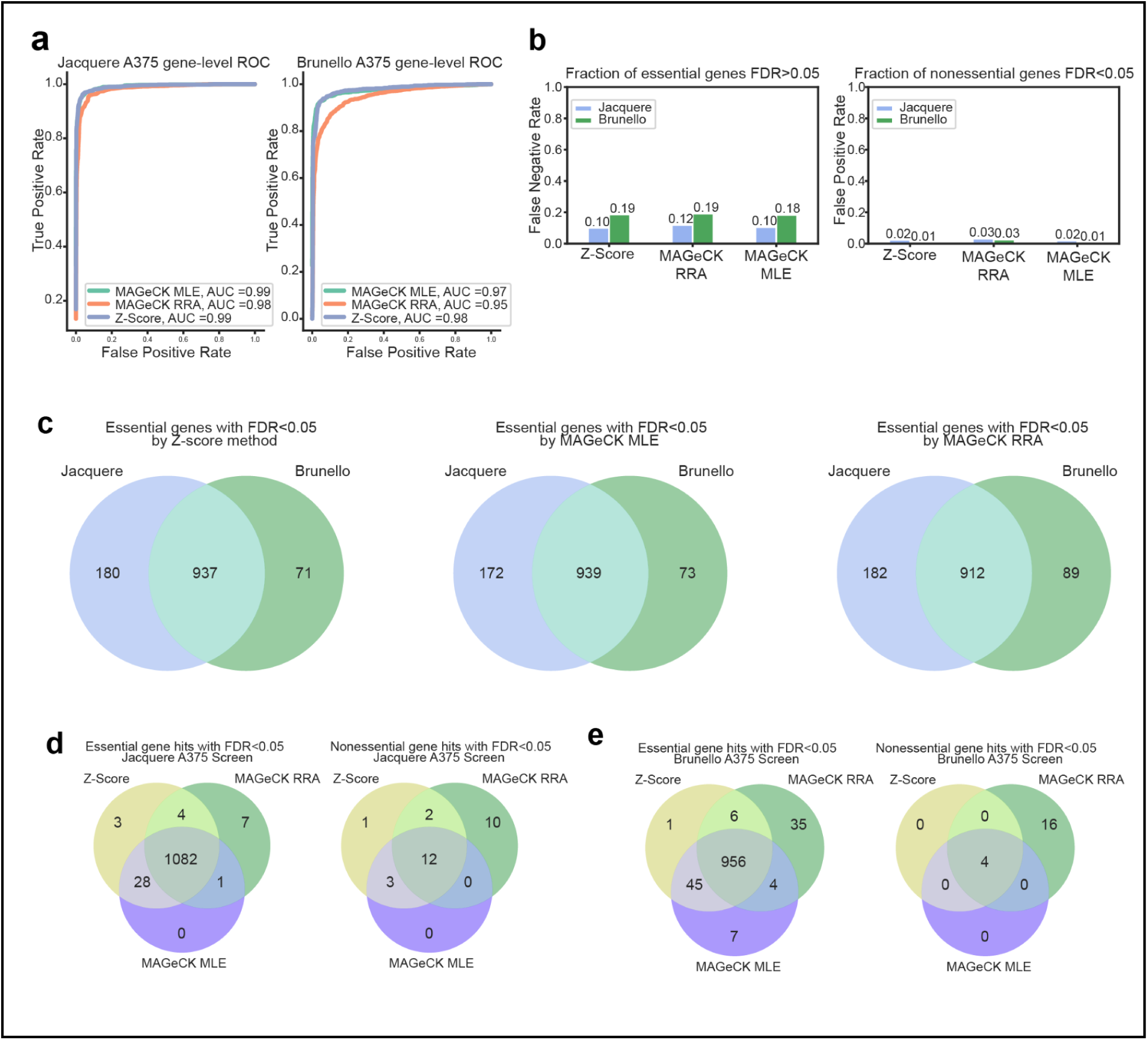
Examination of CRISPR screen hit-calling methods. a) Receiver operating curve evaluating the ability of each hit-calling method to discretize common essential from nonessential genes with screening data of the Brunello and Jacquere library in the A375 cell line. b) Evaluation of the false discovery rate (FDR) threshold of 0.05 to identify genes with a selective effect in Brunello and Jacquere (A375) screening data for each hit-calling method. The false negative rate represents the fraction of common essential genes targeted by the library that fail to emerge as a hit at FDR<0.05, whereas the false positive rate represents the fraction of nonessential genes that emerge as a hit. c) Comparison of gene hits identified at an FDR threshold of <0.05 on Brunello and Jacquere screening data, both in the A375 cell line. Analysis is repeated with the use of the MAGeCK RRA, MAGeCK MLE, and Z-score method. d) Comparison of gene hits identified by MAGeCK RRA, MAGeCK MLE, and the Z-score method on Jacquere library A375 screening data. Venn diagrams represent hits determined by each method (FDR<0.05) for common essential (left), and common nonessential genes (right). e) Comparison of gene hits identified by MAGeCK RRA, MAGeCK MLE, and the Z-score method on Brunello library A375 screening data. Venn diagrams represent hits determined by each method (FDR<0.05) for common essential (left), and common nonessential genes (right).

The divergence in hits identified by both the Z-score method and MAGeCK MLE, which use effect sizes to determine gene effects, and the rank-based MAGeCK RRA method reveals the limitations of the latter approach. True essential genes that MAGeCK RRA alone fails to recognize as hits in the Jacquere screen (n=28) are represented by guides that deplete relative to intergenic targeting guides but individually feature disparate levels of depletion, demonstrating the disadvantage of the model’s assumption that genes with no effect have evenly distributed guide ranks (Supplementary Figure 4a). Additionally, among nonessential genes that only MAGeCK RRA falsely identifies as hits (n=10), several feature only a single guide that is depleted (Supplementary Figure 4b). This result reflects how, in order to determine if a gene is a hit, the method ignores the information offered by all but the most outlying guide targeting a gene when the others are assumed to be ineffective.

The results of the MAGeCK RRA hit-calling method therefore allow genes to be falsely associated with the screening behavior of individual guides that are not representative of target gene function, such as those with substantial off-target activity. Nevertheless, they are interpreted as direct evidence of gene function in practice. For example, a recent study used MAGeCK RRA p-values as evidence to report genes that improve the tumor-infiltrating capability of natural killer cells ^60^. The MAGeCK RRA output file provided in that study reveals that almost none of the positive selection hits reported, including the focal point of the manuscript *Calhm2*, are targeted by any of the top 25% of positively-selected guides in the library. Thus, these genes are determined to have a positive selective effect in the absence of evidence of enrichment from even a single guide. The lack of positive selection among guides targeting the reported hit genes is also evident by direct comparison to the non-targeting controls (Supplementary Figure 5). Indeed, while acknowledging its potential for false negatives, the Z-score method indicates that all reported hits are statistically insignificant (FDR > 0.05).

In summary, we urge large scale perturbation screen studies to be aware of both false positive and false negative modes of any particular analytical framework. Since the strongest screen hits will emerge as significant with any well-reasoned hit-calling method, implementing more than one method and identifying gene hits at the intersection of results is merited. Such an approach will mitigate the reporting of false discoveries that, if left unchecked, would undermine confidence in large-scale perturbational screens^61^.

## DISCUSSION

We are now equipped with state-of-the-art tools to predict rates of Cas9 CRISPR knockout guide activity at target and off-target sites with high precision. However, prioritizing off-target avoidance in library design decreases the representation of effective guides, rendering the resulting screens susceptible to false negatives. One approach to minimize false negatives is to increase the number of guides per gene, but this is restricted by scale, especially for resource-intensive yet highly informative screening methods such as CROP-seq that demand compact screening libraries. We therefore encourage minimizing false negatives in the design of primary screens through conservative avoidance of false positives, which can be later identified with secondary screening or direct guide validation. Here, we demonstrate such an approach that accounts for off-target activity in a data-driven manner while adequately prioritizing selection of guides with strong on-target activity as predicted by the Rule Set 3 model. The resulting library, Jacquere, exhibits satisfactory screening performance and particularly effective control of false negatives when screened in A375 and A549 cell models.

Another potential source of excessive double-stranded breaks in CRISPR knockout screens are copy number amplified regions present in the cell model. The design of Jacquere is cell-line agnostic for the sake of broad application and thus does account for the effect of copy number amplified regions present in any given model. Fortunately, analytical methods such as CRISPRCleanR are available to mitigate this effect in library design ^48,62^. To avoid this potential source of error in screen interpretation, we advise careful analysis of genes with copy number amplification in the cell model employed.

While here we emphasize design considerations with respect to the human genome, the Aggregate CFD metric and guide selection approach introduced in this manuscript is applicable to the design of Cas9 CRISPR knockout screening libraries against other genomes. We designed an updated genome-wide mouse library, Julianna (see Methods), which mirrors the underlying design principles of Jacquere. For the design of such libraries for additional organisms, we recommend the use of the CRISPick web tool. In future work, we aim to apply a similar framework to guide selection for Cas12a libraries, which offer attractive properties for multiplexing^64,65^. At present, Jacquere reflects currently understood patterns of Cas9 CRISPRko on-target and off-target activity, a strategic balance of key criteria in library design, and high coverage of the contemporary, comprehensive set of annotated human protein coding genes in a compact design to empower a wide variety of screening projects.

## METHODS

### Analysis of GUIDE-seq data

GUIDE-seq data from the Yaish and Orenstein study was downloaded from GitHub (github.com/OrensteinLab/CRISPR-Bulge) on April 16, 2024 ^40^. Reads corresponding to each guide were divided by the number of reads at the on-target site within the experiment, then these normalized reads were averaged for guides repeated across experiments. Active off-target sites are defined as those at least 1% as active as the on-target site (>0.01 normalized reads). Among the 5,857,388 potential off-target sites identified by SWOffinder for the 114 guides represented, 1,350 are active by this definition.

The Yaish and Orenstein study reported numerous off-target sites predicted to bind with a DNA or RNA bulge, yet the bulge position lies in PAM-distal positions outside of the SDR. Thus, while reads are reported for “bulge” activity, it is likely that activity at these sites occurred as a result of direct binding. Thus, we restricted bulge off-target sites to those that require a DNA or RNA bulge in order to have fewer than three mismatches in the SDR. The source paper acknowledges the limited precision of the GUIDE-seq technology, stating that the true sites of off-target activity may lie anywhere in a 73 nucleotide window within a sequencing read. We therefore evaluated all possible alignments along the site window for each of the off-target sites to obtain the minimum mismatch count within the SDR.

### Quantifying computational complexity associated with off-target guide evaluation

The reported computational complexity of the search for off-target sites reflects the number of queries against the genome required by the CRISPick web tool. This number is determined by the equation 3^m^ ✕ *choose*(l,m), where m represents the maximum tolerated number of mismatches, and l represents the length of the guide sequence in consideration (such as 19 for the specificity-defining region introduced in this manuscript). The total count of candidate guides for a genome-wide CRISPR screen represents all sites with the *Streptococcus pyogenes* Cas9 canonical PAM NGG along the protein coding region of all Ensembl v113 genes as reported in the CRISPick web tool genome-scale guide designs.

### CRISPick Aggregate CFD training data cleaning

The training dataset is published in DeWeirdt et al. (2022) and employs SpCas9 CRISPRko with the Chen et al. tracrRNA in A375 Cells ^27,66^. The complete tiling library includes 48,742 guides to target 201 essential genes, 33,643 guides to target 198 nonessential genes, 1000 intergenic controls, and 1000 non-targeting controls. The essential and nonessential genes were randomly selected from gold standard gene lists ^67,68^. Here, guide sequences were mapped to Ensembl v113 annotations to determine the target gene. Raw count matrices were log-normalized and filtered to exclude guides with low representation, as indicated by pDNA counts three or more standard deviations below the mean. Log fold change in guide abundance was calculated relative to the pDNA. Biological replicates were averaged upon confirming reproducibility (Pearson correlation of log fold changes 0.77, 0.70, and 0.79 between replicates A, B, and C). Z-scores for each guide were calculated using the standard deviation and mean of intergenic control log fold changes, which represent the expected depletion associated with a single double-stranded break.

To identify nonessential-targeting guides, the data was subset to those targeting genes that have previously been identified as nonessential ^68^. Genes with an average z-score relative to the intergenic controls of below −2 were omitted from the previously-defined set of nonessential genes since they are understood to have an essential function or copy number amplification in this cell line. Guides with a z-score below −3 were defined as promiscuous and above −1 as non-promiscuous, thereby excluding guides with ambiguous classification from the training set. Upon recognition that inactive guides would introduce noise in classifier training given their tendency to not generate DSBs regardless of predicted off-target sites, the data was restricted to effective guides as determined by their sequence composition (RS3 Sequence Score >0.2). The final training set consists of 685 promiscuous guides and 7,166 non-promiscuous guides targeting 165 nonessential genes.

### CRISPick Aggregate CFD validation data cleaning

Screening data of the Avana Library was downloaded from DepMap 24Q2 ^19^. To identify the extent to which Aggregate CFD performance depends on a cell’s degree of sensitivity to double-stranded breaks, cell lines were separated by TP53 mutation status as classified by prior efforts, where TP53 wildtype cell lines represent those more prone to an anti-proliferative response in the presence of double-stranded breaks ^48^. Within each set, each guide is represented by the median count across cell lines. As in the analysis of Deweirdt 2022 tiling data, nonessential-targeting genes were identified from the gold standard set and filtered to exclude those with depletion across guides, resulting in 781 nonessential genes. Since this data does not include intergenic controls, promiscuous nonessential-targeting guides were identified as those with a log fold change at least two standard deviations below the nonessential-targeting mean (n=108, 116 for TP53 mutant, wildtype cells). Non-promiscuous guides were identified as those comparable to the non-targeting controls, namely those with log-fold changes at least within one standard deviation of the mean of non-targeting controls (n=1550, 1355 for TP53 mutant, wildtype cells).

### Jacquere Library Design

The Jacquere library was designed against the human genome (GRCh38) and targeted the comprehensive set of protein-coding genes annotated by RefSeq (version 08-2024), GENCODE (version 47), and CHESS 3.1.3. The set of protein coding genes annotated by RefSeq and GENCODE was obtained by running CRISPick with NCBI and Ensembl as the reference genomes respectively and retrieving the unique set of Target Gene Symbols from the design results. Protein coding genes annotated by the CHESS database were obtained from the file chess3.1.3.GRCh38.assembly.gtf on the CHESS downloads page (ccb.jhu.edu/chess/). The three resulting gene lists were then pasted into the HGNC database to obtain approved gene symbols and identify genes that are identical despite distinct IDs across the aforementioned sources ^69^. Lastly, a custom python script was used to identify the overlap and discrepancies between databases.

The CRISPick web tool was employed for the design of the Jacquere library, specified to *Streptococcus pyogenes* Cas9, a quota of four, and Target Local Mode. For each target gene, CRISPick identifies the full set of candidate guides as those with the canonical PAM site on either strand of the target, excludes those containing BsmBI restriction sites or with a run of four or more thymidines. Guides for each target are then selected by order of their RS3 score (sequence+target, optimized for use of the Chen tracrRNA), relaxing key guide criteria as required to fill the target-level quota (Supplementary Data 1). In order to target genes annotated across databases, Jacquere was designed with 4 CRISPick runs:

1. Targets set of all protein-coding genes with Ensembl (v113) as reference genome to target all annotated by GENCODE (v47)
2. Targets set of all protein-coding genes that are annotated by RefSeq (v2024_08) but not GENCODE, using NCBI as the reference genome
3. Targets genes annotated uniquely by CHESS (v3.1.3) by supplying the exon coordinates of the primary CHESS transcripts. We use Ensembl v113 as the reference genome, which determines the classification of guide off-target sites.
4. Generate 100 non-targeting controls in CRISPick

Lastly, intergenic controls for Jacquere were designed by generating a set of 10,000 possible intergenic control sequences, retrieving their coordinates using the UCSC BLAT command line tool, and supplying the coordinates as targets for CRISPick to retrieve Aggregate CFD and RS3 scores. Intergenics were then subsampled to 900 that resemble the distribution of CRISPick Aggregate CFD and RS3 scores to that of targeting guides in Jacquere, allowing them to serve as representative controls (Supplementary Figure 2b).

After concatenating the CRISPick runs, the library was filtered to exclude guides that exceed a CRISPick Aggregate CFD of 4.8 and subset to the top 3 guides per gene. The gene-level pick orders from CRISPick are provided to support the use of this library with a reduced or increased quota (Supplementary Data 4). 166 of the genes identified across RefSeq, GENCODE, and CHESS are not targetable by the criteria we implement in the design of Jacquere (Supplementary Data 5). Such genes are only targeted in prior libraries by guides that exceed a CRISPick Aggregate CFD of 4.8 or indicate severely low potential for on-target activity by violating multiple guide selection criteria.

The CRISPick web tool targets unique genes on the basis of their Gene ID rather than Gene Symbol. There are 27 gene symbols that are associated with more than one unique gene, as distinguished by Ensembl gene ID, and the Jacquere designs demonstrate that 80% of these genes are optimally targeted by a distinct set of guides from other genes that share the same symbol.

### Julianna Library Design

We provide designs for Julianna, a novel Cas9 CRISPRko genome-wide library targeting the mouse genome (Supplementary Data 6). The Julianna Library implements the same guide selection approach as Jacquere, only eliminating restrictions on target site variability due to the absence of population-level mouse data. To design Julianna, genome-wide Cas9 CRISPRko guide selections against the GRCm38 build and Ensembl v102 annotations were obtained with CRISPick. As in Jacquere, the CRISPick run targeted the complete set of protein coding genes using a quota of 4 and Target Local Mode. Guides with a CRISPick Aggregate CFD exceeding 4.8 were manually removed from the library. The standard Julianna library features a quota of 3, yet we additionally provide designs for up to four guides per gene (Supplementary Data 7).

### Assessment of external Cas9 CRISPRko libraries

The GeCKO v2, TKOv3, and Gattinara libraries were downloaded from Addgene, with GeCKO v2 libraries A and B concatenated prior to analysis. The Avana library was downloaded from DepMap (18Q1), and the Brunello and MinLibCas9 libraries were retrieved from their source manuscripts ^15,19,25^. The Project Score library (referred to as KosukeYusa or KY v1 in external works) as reported in its source manuscript indicates 19 nt sequences, which are insufficient for our internal mapping tool. Thus, 20 nt Project Score guide sequences were obtained from the MinLibCas9 project (github.com/EmanuelGoncalves/crispy/blob/master/notebooks/minlib/libraries/MasterLib_v1.csv .gz), selecting guides marked as “KosukeYusa” and removing the 3nt PAM sequence from the end. Lastly, the top 6 genome-wide guide picks per gene according to the VBC score project were downloaded from the online prediction tool (vbc-score.org/download).

Guide mappings to genes for each library were obtained through use of our internal tool to identify the target site in the framework of GENCODE47 annotations, and guides were evaluated for on-target and off-target properties with the CRISPick web tool. Since CRISPick does not evaluate any guides with over 20,000 predicted off-target sites, there are 66, 13, and 7 guides from GeCKO v2, Avana, and Brunello respectively omitted from the counts of guides that violate the aforementioned filters that pertain to off-target activity (Table 1).

False negative rates across Cas9 CRISPRko genome-wide libraries were estimated using each library’s guide selections for the 201 essential-targeting genes in the RS3 validation tiling screen and their respective z-scores (calculated as described in the CRISPick Aggregate CFD training data cleaning section) ^27^. Importantly, the measured activity of these guides was not used in the training of Rule Set 3. For the purpose of this analysis, gene effects were determined by averaging the targeting guides z-scores so that effect sizes are not influenced by set size, thereby enabling unbiased comparison across libraries. The false negative rate of each library was determined from the number of genes that fail to deplete, implementing various effect size thresholds to identify a gene as depleted (Figure 3d). Since this analysis employs a set-size agnostic metric to determine gene effects, it should not be interpreted as evidence of statistical confidence offered by libraries with varying quotas, but rather the ability of such libraries to reflect the true phenotype with the guides employed.

### Jacquere screen experimental methods

#### Table of Reagents

| Reagent or Resource | Source | Identifier |
| --- | --- | --- |
| **Chemicals** |  |  |
| Puromycin | Sigma-Aldrich | Cat#P9620 |
| Blasticidin | Thermo Fisher | Cat#A1113903 |
| Polybrene | Sigma-Aldrich | Cat#H9268 |
| TransIT-LT1 | Mirus | Cat#2306 |
| **Experimental models: Cell lines** |  |  |
| Human A549 cells | Cancer Cell Line Encyclopedia | N/A |
| Human A375 cells | Cancer Cell Line Encyclopedia | N/A |
| Human HEK293T cells | ATCC | ATCC CRL-3216 |
| **Recombinant DNA** |  |  |
| Plasmid pMV_AA088 | Addgene | Addgene #216104 |
| Plasmid pMV_AA050 | Addgene | Addgene #216107 |
| Plasmid pRDA_734 | Addgene | Addgene #216097 |
| **Software and algorithms** |  |  |
| PoolQ | The Broad Institute | https://portals.broadinstitute.org/gpp/public/software/poolq |

#### Modular vector design

All pF plasmids were made by gene synthesis into the EcoRV site of pUC57-Kan (Genscript).

All pF vectors were assembled into plasmids using Golden Gate cloning.

pMV_AA088 (Addgene #216104): EF1α promoter expresses SpyoCas9 and blasticidin resistance; split by a 2A site.

pMV_AA050 (Addgene #216107): U6 promoter expresses sgRNA (library cloned via BsmBI sites); EF1α promoter expresses puromycin resistance.

pRDA_734 (Addgene #216097): U6 promoter expresses sgRNA (library cloned via BsmBI sites); EFS promoter expresses SpyoCas9 and puromycin resistance; split by a 2A site.

#### Modular vector destination vector pre-digest

Destination vectors (pRDA_734, pMV_AA050) were pre-digested using BbsI and NEBuffer 2.0 (New England Biolabs) at 37°C for 2 h. Linearized destination vectors were gel purified using 0.7% agarose gels and extracted with the Monarch DNA Gel Extraction Kit (New England Biolabs), before further purification by isopropanol precipitation.

#### Modular vector Golden Gate assembly

The pF vectors were diluted to 10 nM in sterile water and cloned into a pre-digested destination vector via Golden Gate cloning. Each reaction contained 3 μL BbsI (New England Biolabs), 1.25 μL T4 ligase (New England Biolabs), 3 μL of 10x T4 ligase buffer (New England Biolabs), 75 ng destination vector, and a 1:1 molar ratio of fragments:destination vector. Reactions were carried out under the following thermocycler conditions: (1) 37°C for 5 min; (2) 16°C for 5 min; (3) go to (1), x100; (4) 37°C for 30 min; (5) 65°C for 20 min. The Golden Gate product was treated with Exonuclease V (New England Biolabs) at 37°C for 30 min before enzyme inactivation with the addition of EDTA to 11 mM. Per reaction, 10 μL of product was transformed into Stbl3 chemically competent E. coli (Invitrogen) via heat shock, and grown at 37°C for 16 h on agar with 100 μg/mL carbenicillin. Colonies were picked and grown at 37°C for 16 h in 5 mL Luria-Bertani (LB) broth with 100 μg/mL carbenicillin. Plasmid DNA (pDNA) was prepared (QIAprep Spin Miniprep Kit, Qiagen). Purified plasmids were verified by restriction enzyme digest and whole plasmid sequencing through Plasmidsaurus.

#### Library production

Oligonucleotide pools were synthesized by Twist. BsmBI recognition sites were appended to each guides sequence along with the appropriate forward and reverse overhang sequences (bold italic) for cloning into the sgRNA expression plasmids, as well as primer sites to allow differential amplification of subsets from the same synthesis pool. The final oligonucleotide sequence structure was thus: 5′-[Forward Primer]CGTCTCA***CACC***G[guide, 20 nt]***GTTT***CGAGACG[Reverse Primer]-3’.

Primers were used to amplify individual subpools using 25 μL 2x NEBnext PCR master mix (New England Biolabs), 1-2 μL of oligonucleotide pool (∼30-300 ng), 5 μL of primer mix at a final concentration of 0.5 μM, and water to a final volume of 50 µL. PCR cycling conditions: (1) 98°C for 1 min; (2) 98°C for 30 sec; (3) 53°C for 30 s; (4) 72°C for 30 s; (5) go to step 2, x6-14 depending on library size; (6) 72°C for 5 min.

The resulting amplicons were PCR-purified (Qiagen) and cloned into their respective library vector via Golden Gate cloning with Esp3I (Fisher Scientific) and T7 ligase (Epizyme) under the following thermocycler conditions: (1) 37°C for 5 min; (2) 20°C for 5 min; (3) go to step 1, x99; (4) 37°C for 30 min; (5) 65°C for 10 min. The ligated product was isopropanol precipitated and electroporated into Stbl4 electrocompetent cells (Invitrogen) and grown at 37°C for 18 h on agar with 100 μg/mL carbenicillin. Colonies were scraped and plasmid DNA (pDNA) was prepared (HiSpeed Plasmid Maxi, Qiagen). To confirm library representation and distribution, the pDNA was sequenced by Illumina MiSeq.

#### Cell culture

All cells regularly tested negative for mycoplasma contamination and were maintained in the absence of antibiotics except during screens and lentivirus production, during which media was supplemented with 1% penicillin–streptomycin. Cells were passaged every 2–3 days to maintain exponential growth and were kept in a humidity-controlled 37 °C incubator with 5.0% CO2. A549 cells were passaged with DMEM supplemented with 10% Fetal Bovine Serum (FBS). A375 cells were passaged with RPMI supplemented with 10% FBS. HEK293T cells were passaged with DMEM supplemented with 10% heat-inactivated FBS.

#### Lentivirus production

For small-scale virus production, the following procedure was used: 18 h before transfection, HEK293T cells were seeded in 6-well dishes at a density of 1×10^6^ cells per well in 1 mL of DMEM + 10% heat-inactivated FBS. Transfection was performed using the TransIT-LT1 transfection reagent (Mirus) according to the manufacturer’s protocol. Briefly, for each construct, 324μL of Opti-MEM (Corning) and 17μL LT1 was combined with a DNA mixture of the packaging plasmid pCMV_VSVG (125 ng; Addgene 8454), psPAX2 (1250 ng; Addgene 12260), and the transfer vector (312.5 ng). The solutions were incubated at room temperature for 30 min and added dropwise to cells. Plates were then transferred to a 37°C incubator for 6–8 h, after which the media was removed and replaced with DMEM + 10% FBS media supplemented with 1% BSA. Virus was harvested and filtered 40 hours after this media change.

A larger-scale procedure was used for pooled library production. 18 h before transfection, 18×10^6^ HEK293T cells were seeded in a 175 cm^2^ tissue culture flask and the transfection was performed the same as for small-scale production using 6 mL of Opti-MEM, 305 μL of LT1, and a DNA mixture of pCMV_VSVG (5 μg), psPAX2 (50 μg), and 10 μg of the transfer vector. Following addition of the transfection mix, flasks were transferred to a 37°C incubator for 6–8 h, then the media was aspirated and replaced with BSA-supplemented media; virus was harvested and filtered 40 h after this media change.

#### Determination of lentiviral titer

To determine lentiviral titer for transductions, cell lines were transduced in 12-well plates with a range of virus volumes (e.g., 0, 150, 300, 500, and 800 μL virus) with either 1.5×10^6^ or 3×10^6^ cells per well in the presence of polybrene. Larger constructs, such as the all-in-one vector pRDA_734, used 1.5×10^6^ cells per well. For transduction, the plates were centrifuged at 821 x g for 2 h, after which 2 mL of warm media was added to reduce viral toxicity. Plates were then transferred to a 37°C incubator for 4–6 h. Each well was then trypsinized and pooled. Two days post-transduction, we seeded an equal number of cells into two wells of a 6-well plate, and added puromycin to one well. The following passage, both wells were counted for viability. A viral dose resulting in ∼30% transduction efficiency, corresponding to an MOI of ∼0.35, was used for subsequent library screening.

#### Small molecule dosages

The dosages for the selection drugs puromycin and blasticidin were as follows for the relevant cell lines:

A549: puromycin 1.5 μg/mL; blasticidin 5 μg/mL

A375: puromycin 1 μg/mL; blasticidin 5 μg/mL

Puromycin selection was completed over 5-7 days, while blasticidin selection was completed over 12-14 days.

#### Lentiviral transduction to establish stable cell lines

In order to establish stable Cas-expressing cell lines for split-vector screens, A375 cells or A549 cells were transduced with pMV_AA088 in the presence of polybrene at a dosage of 1 µg/mL. Cells were centrifuged at 821 x g for 2 h in 12-well plates, after which 2 mL of warm media was added to reduce viral toxicity. Plates were then transferred to a 37°C incubator for 4–6 h.

Replicate wells were then trypsinized and pooled. Successfully infected cells were selected for with blasticidin as described above.

#### Pooled screens

For pooled screens, cells were transduced in 2 biological replicates with a lentiviral library. Transductions were performed at a low multiplicity of infection (MOI ∼0.35), using enough cells to achieve a representation of at least 500 transduced cells per guide assuming a 20% - 40% transduction efficiency. The transduction protocol was the same as listed above. Puromycin was added 2 days post-transduction. Cells were passaged on puromycin for 2 passages to ensure complete removal of non-transduced cells. After selection, cells were passaged at at least 500x representation every 2-3 days for an additional 2 weeks to allow guides to enrich or deplete; cell counts were taken at each passage to monitor growth. At the conclusion of each screen (Day 21 after transduction of the guide library), cells were pelleted by centrifugation, resuspended in PBS, and frozen promptly for genomic DNA isolation.

#### Genomic DNA isolation, PCR, and sequencing

Genomic DNA (gDNA) was isolated using either the KingFisher Flex Purification System with the Mag-Bind Blood & Tissue DNA HDQ Kit (Omega Bio-Tek), per the manufacturer’s instructions. The gDNA concentrations were measured by Qubit.

For PCR amplification, unless otherwise noted, gDNA was divided into 100 µL reactions such that each well had at most 10 µg of gDNA. Plasmid DNA (pDNA) was also included at a maximum of 100 pg per well. Each well of a 96-well PCR plate contained 1.5 µL of Titanium Taq (Takara), 10 µL of Titanium Taq buffer, 8 µL of dNTP Mix, 5 µL of DMSO, 0.5 µL of P5 primer at 100 µM stock, 10 µL of P7 primer at 5µM stock, and nuclease-free water added to 100 µL. PCR cycling conditions were as follows: (1) 95°C for 1 min; (2) 94°C for 30 s, (3) 52°C for 30 s, (4) 72°C for 30 s, (5) go to step 1, x27; (6) 72°C for 10 min. PCR products were pooled and purified with Agencourt AMPure XP SPRI beads according to manufacturer’s instructions (Beckman Coulter, A63880). Samples were sequenced using Illumina technology NovaSeq 6000 100 cycles, single-read, with a 5% spike-in of PhiX.

Guide sequences were extracted from sequencing reads by running PoolQ with the search prefix “CACCG”, and reads were counted by alignment to a reference file of all possible guides present in the library.

##### Analysis of Jacquere and Brunello screening data

Following deconvolution, the resulting matrix of read counts was normalized to reads per million by the following formula: read per guide/total reads per condition×10^6^. Normalized counts were log2-transformed, and the Day 21 (post-transduction) values were subtracted from the plasmid DNA (pDNA) to obtain the log2 fold change for each guide. Log2 fold changes for each guide were averaged across biological replicates upon confirmation of reproducibility (Figure 4a, Supplementary Figure 3a).

Guide level z-score was calculated from log2 fold changes using the standard deviation and mean of intergenic controls, which represent the expected depletion associated with a single double-stranded break. Guides were associated with all genes to which they map, inclusive of all annotations recognized across RefSeq 08-2024 and GENCODE 47, rather than the intended target genes to exemplify the use of current and accurate gene annotations for downstream analysis as recommended in prior works ^26^. Guide-level z-scores were aggregated on a gene level using Stouffer’s method. Since the intergenic guides are not normally distributed (D’Agostino-Pearson K^2^, p<0.01), an expected result of the intergenic guides being designed to match the Aggregate CFD distribution of targeting guides in the Jacquere library, neither the guide nor gene-level z-scores should not be directly interpreted as evidence of statistical significance. Instead, we determine significance by testing the null hypothesis that a gene has the same effect as intergenic-targeting guides, which reflect the phenotype of cells that have sustained a double-stranded break but no loss of function in any gene. This is done by sampling groups of intergenic guides to match the quota of targeting guides in the library, aggregating the guides in the resulting “pseudogene” with Stouffer’s method, smoothing the pseudogene distribution with kernel density estimation, and calculating the percentage of the distribution with a more extreme z-score score than each target gene to yield empirical p-values. False discovery rates are then calculated with the Benjamini-Hochberg correction for multiple testing.

The Brunello A375 screen read counts were downloaded from Supplementary Table 1 of Sanson et al. (2018), using the “A375_mod_tracr raw reads” for consistency with the tracrRNA employed in the Jacquere screen ^50^. The same normalization, log2-transformation, and fold-change calculation as the Jacquere screen was applied, and log2 fold changes were averaged across biological replicates. Guide level-z-scores were calculated from intergenic controls, which are normally distributed (D’Agostino-Pearson K^2^, p=0.275). Guides were associated with all genes to which they map in GENCODE 47 annotations, and gene-level z-scores were determined by aggregating the z-scores of all mapping guides with Stouffer’s method.

To identify library performance, true positive and negative classifications were determined from previously established lists of essential and nonessential genes ^67,68^.

#### Comparison of hit-calling methods

For comparison to the Z-score method implemented in the prior section, MAGeCK RRA and MAGeCK MLE were applied to the Jacquere A375 (pRDA_734 vector) and Brunello A375 screening data ^50^ using the same guide mappings. MAGeCK version 0.5.9.4 was implemented on the command line with the following usage for MAGeCK RRA, exemplified with the Jacquere screening data. To note, the normalization method is specified to “total” instead of the default “median” to remain consistent with the normalization approach we employ for the Z-Score method. This procedural step improves the odds that discrepancies between the results of these methods reflect their distinct approaches to aggregate guides for each gene.

mageck test -k mageck_input_jacquere_A375counts.txt -t

A375_CP0082_repA,A375_CP0082_repB -c CP0082_pDNA -n jacquere --control-sgrna

mageck_input_jacquere_controls_A375.txt --norm-method total

MAGeCK MLE was implemented with the following specifications. The --no-permutation-by-group flag instructs the method to skip calculation of empirical p-values; the permutation test in MAGeCK MLE constructs the null distribution from the set of all guides, thereby rendering empirical p-values subject to the library composition. This study only uses the beta scores from the MAGeCK MLE output to allow pseudogenes constructed from intergenic-targeting guides to serve as the null distribution, so the permutation test is omitted to decrease running time.

mageck mle -k mageck_input_jacquere_A375counts.txt --day0-label CP0082_pDNA

--control-sgrna mageck_input_jacquere_controls_A375.txt --norm-method total

--no-permutation-by-group

MAGeCK MLE exhibits a tendency to excessively identify hit genes as those with fewer than three guides per gene; genes below quota comprise 66.7% of hit genes identified uniquely by MAGeCK MLE upon the Jacquere data in contrast to 0.3% for MAGeCK RRA and 6.5% for the Z-score method. Thus, to generally inform the relative performance of these methods on a three-guide or four-guide per gene library, this study was restricted to genes with guide coverage at the intended quota.

## Code availability

The GitHub (github.com/ldrepano/GPP-Jacquere) features the code and source data required to generate all figures and analyses reported in the manuscript.

## ACKNOWLEDGEMENTS

We thank the Genetic Perturbation Platform, especially Thomas Green and Mark Tomko for developing the infrastructure to enable the analyses described in this manuscript and integrating the novel library design scheme into our web tool, CRISPick. We additionally thank Mudra Hegde for her early work in the development of the CRISPick Aggregate CFD metric and Emanuel Gonçalves for a close reading of the manuscript. This work was supported in part by the Functional Genomics Consortium.

---

**Supplementary Figure 1:**
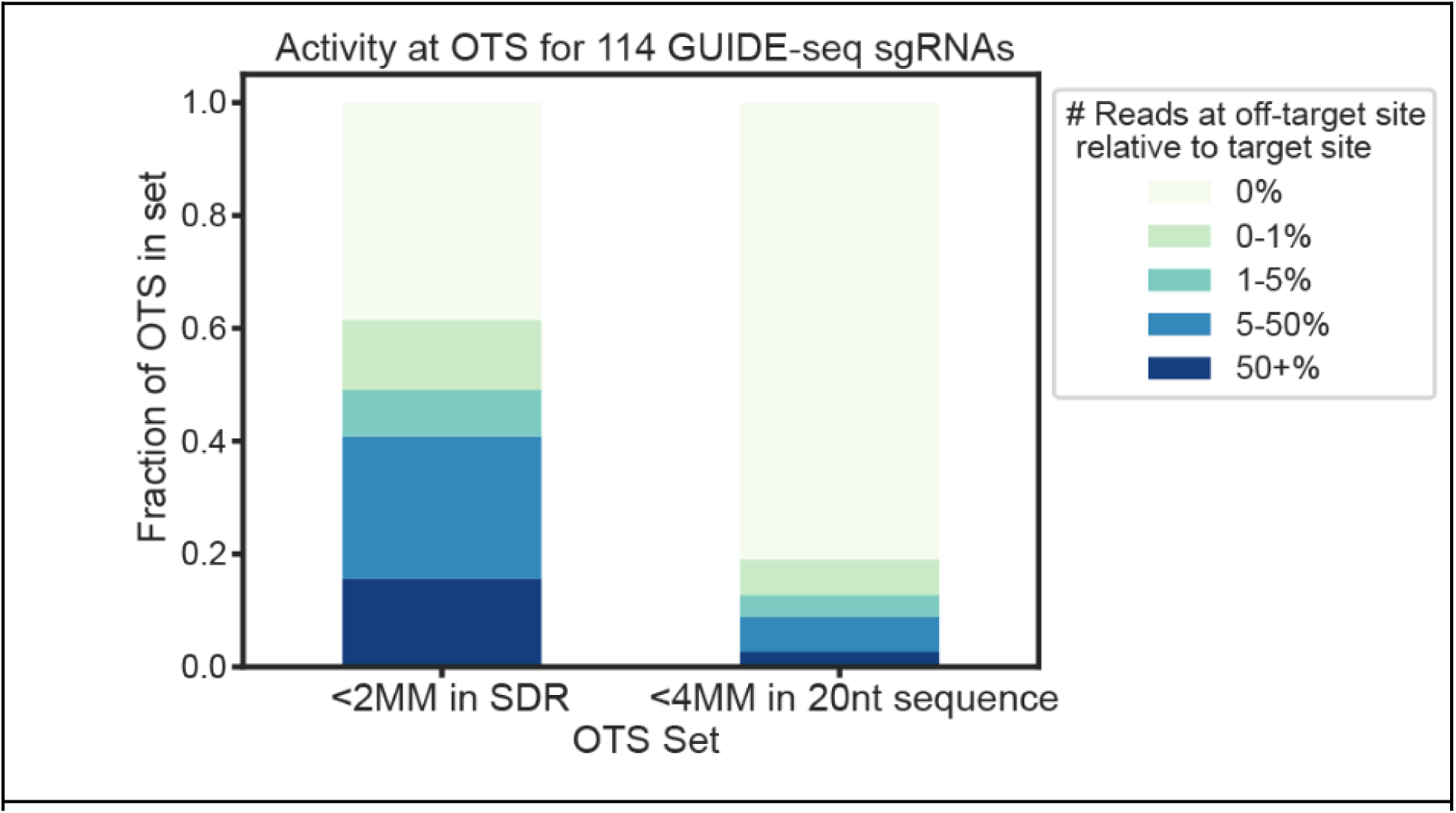
False positive rate corresponding with distinct sets of putative off-target sites as revealed by GUIDE-seq data. GUIDE-seq activity rates of off-target sites (OTS) for the 114 guides in Yaish & Orenstein data, stratified by the set of OTS considered; OTS with up to two mismatches to the guide in the SDR, or OTS with up to four mismatches in the entire 20 nucleotide guide sequence.

**Supplementary Figure 2:**
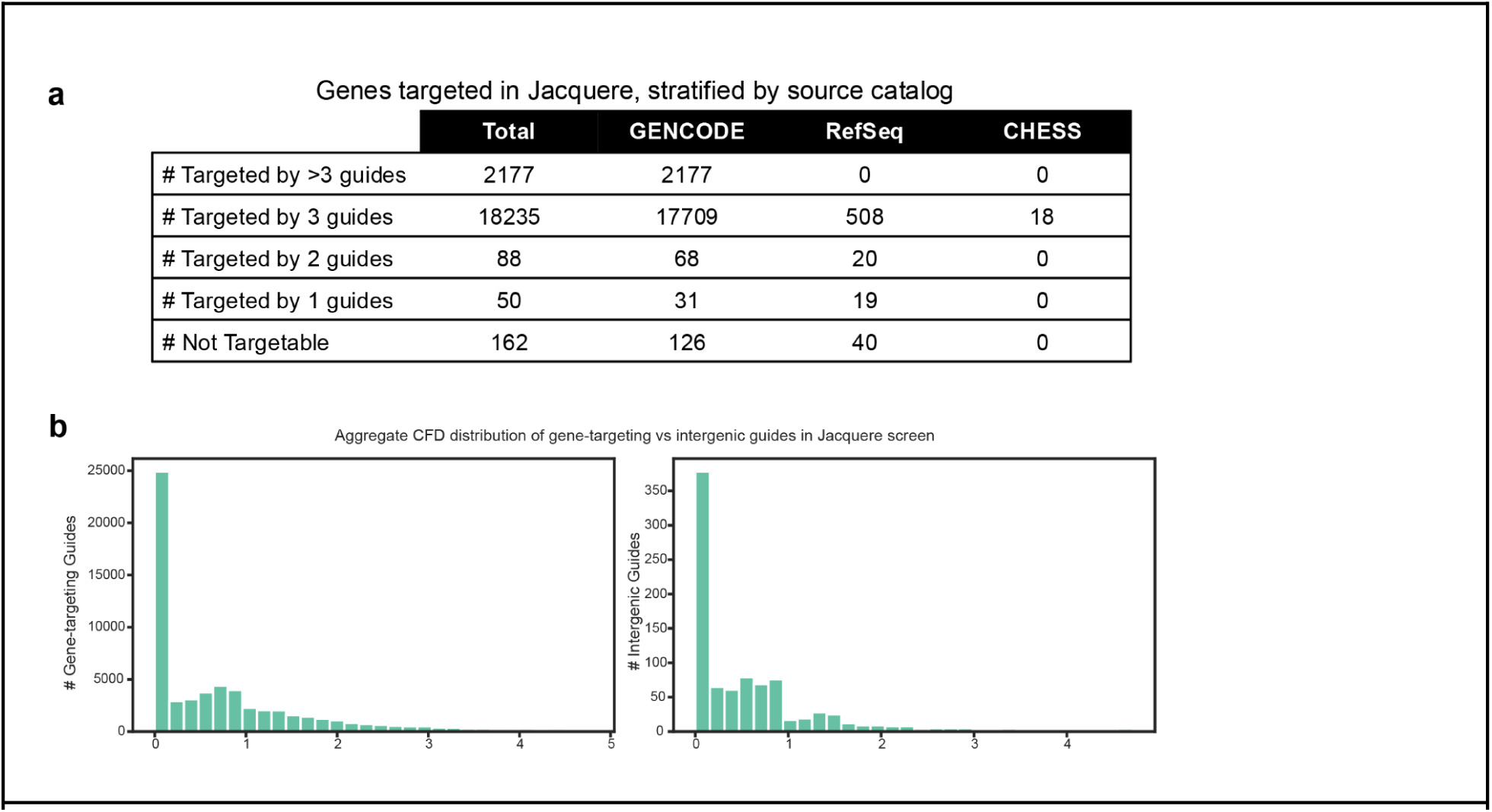
Additional details on the composition of the Jacquere Library. a) The coverage of genes targeted in Jacquere, stratified by whether the gene is present in GENCODE, RefSeq but not GENCODE, or uniquely in CHESS. Genes with incomplete coverage have fewer than three candidate guides with a CRISPick Aggregate CFD of 4.8 or below. Genes are uniquely identified by their Gene ID rather than symbol. b) Distribution of CRISPick Aggregate CFD of gene-targeting and intergenic guides in Jacquere. The average CRISPick Aggregate CFD of intergenic and targeting guides are 0.788 and 0.781.

**Supplementary Figure 3:**
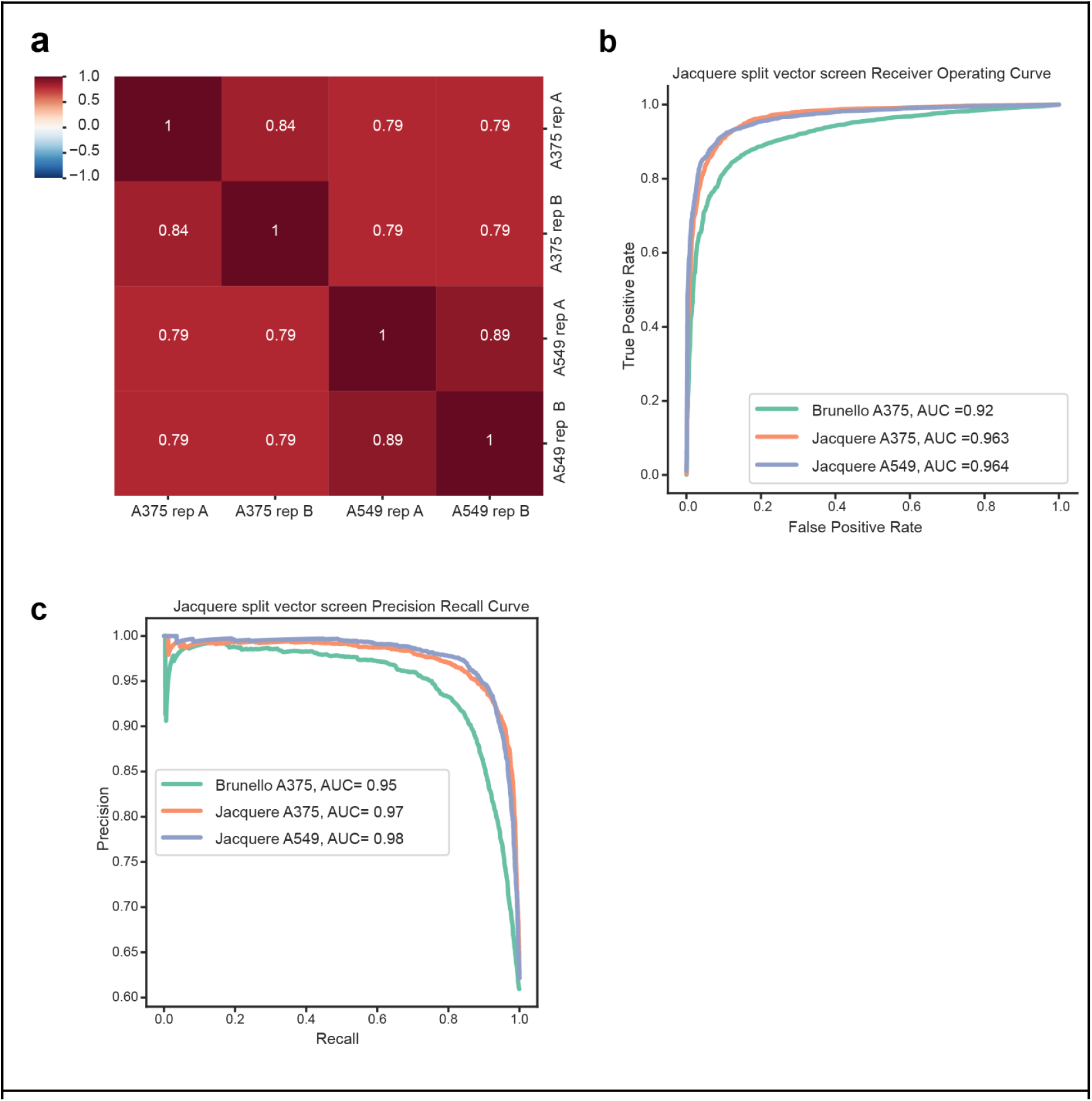
Assessment of Jacquere Screening Performance with separate delivery of the guide and Cas9. a) The Pearson correlation between biological replicates and across cell lines for the Jacquere Library screened with the pMV_AA088 and pMV_AA050 vector. Correlation is calculated upon the log-fold change in guide abundance at Day 21 post-transduction relative to pDNA abundance. b) Receiver operating characteristic analysis of cell viability screening data for the Jacquere library (with the use of the pMV_AA088 and pMV_AA050 vector) and Brunello Library screened in A375 cells. False positive rates are determined by non-essential genes and plotted against the true positive rate, determined by essential genes. c) Precision recall curve to compare the classification of essential genes between Jacquere split vector and Brunello screening data.

**Supplementary Figure 4:**
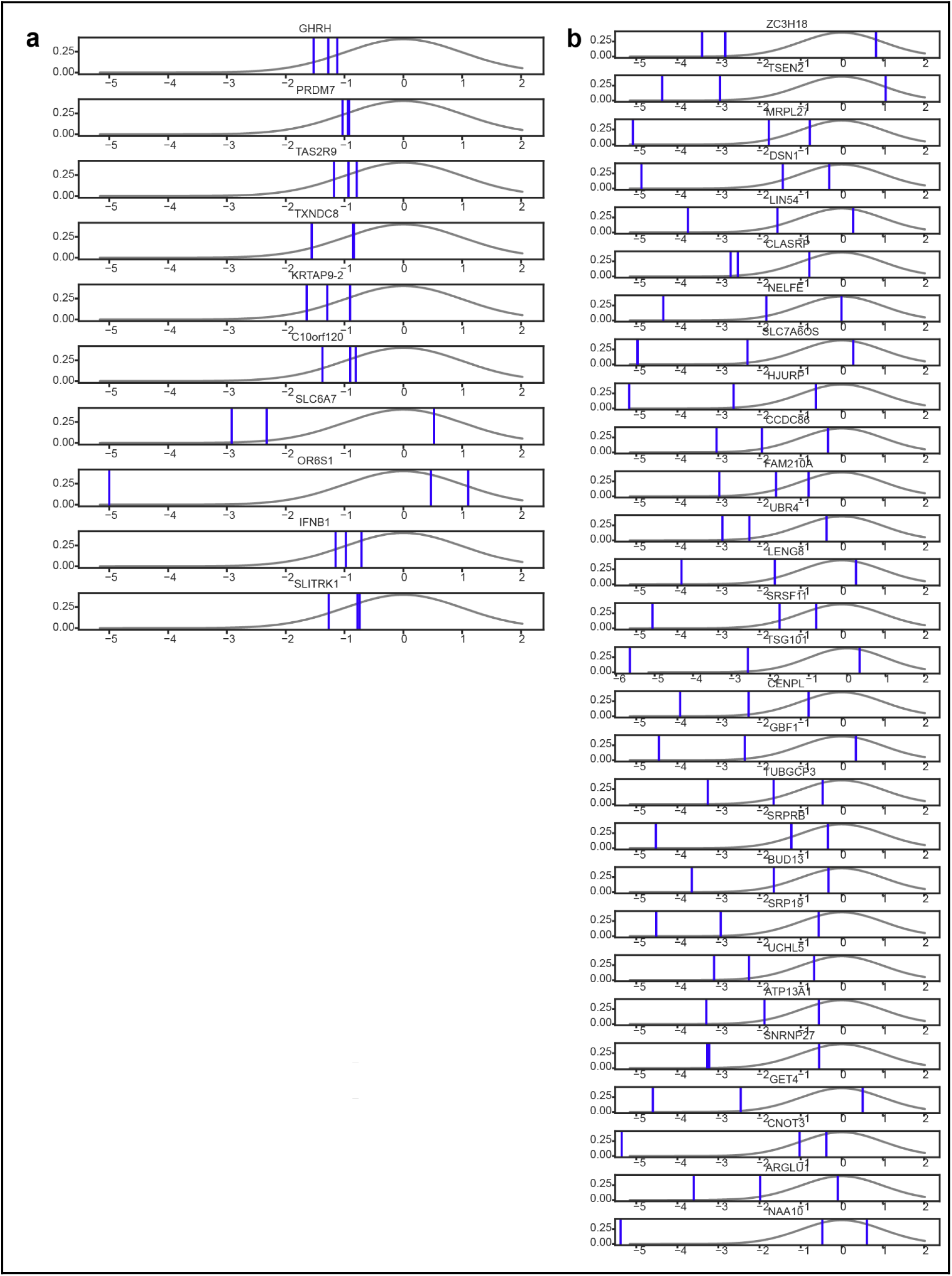
Limitations of hit identification with the MAGeCK RRA method. a) The guide-level behavior of essential genes that are identified as hits by the Z-Score method and MAGeCK MLE but not MAGeCK RRA, thereby representing false negatives that emerge by determining screen hits with this method. The blue lines indicate the z-scores of all guides targeting the gene in the Jacquere library, whereas the background distribution (gray) represents the intergenic-targeting guides. b)The guide-level behavior of nonessential genes that are identified only by the MAGeCK RRA method as hits, thus serving as false positives. The blue lines indicate the z-scores of all guides targeting the gene in the Jacquere library, whereas the background distribution (gray) represents the intergenic-targeting guides.

**Supplementary Figure 5:**
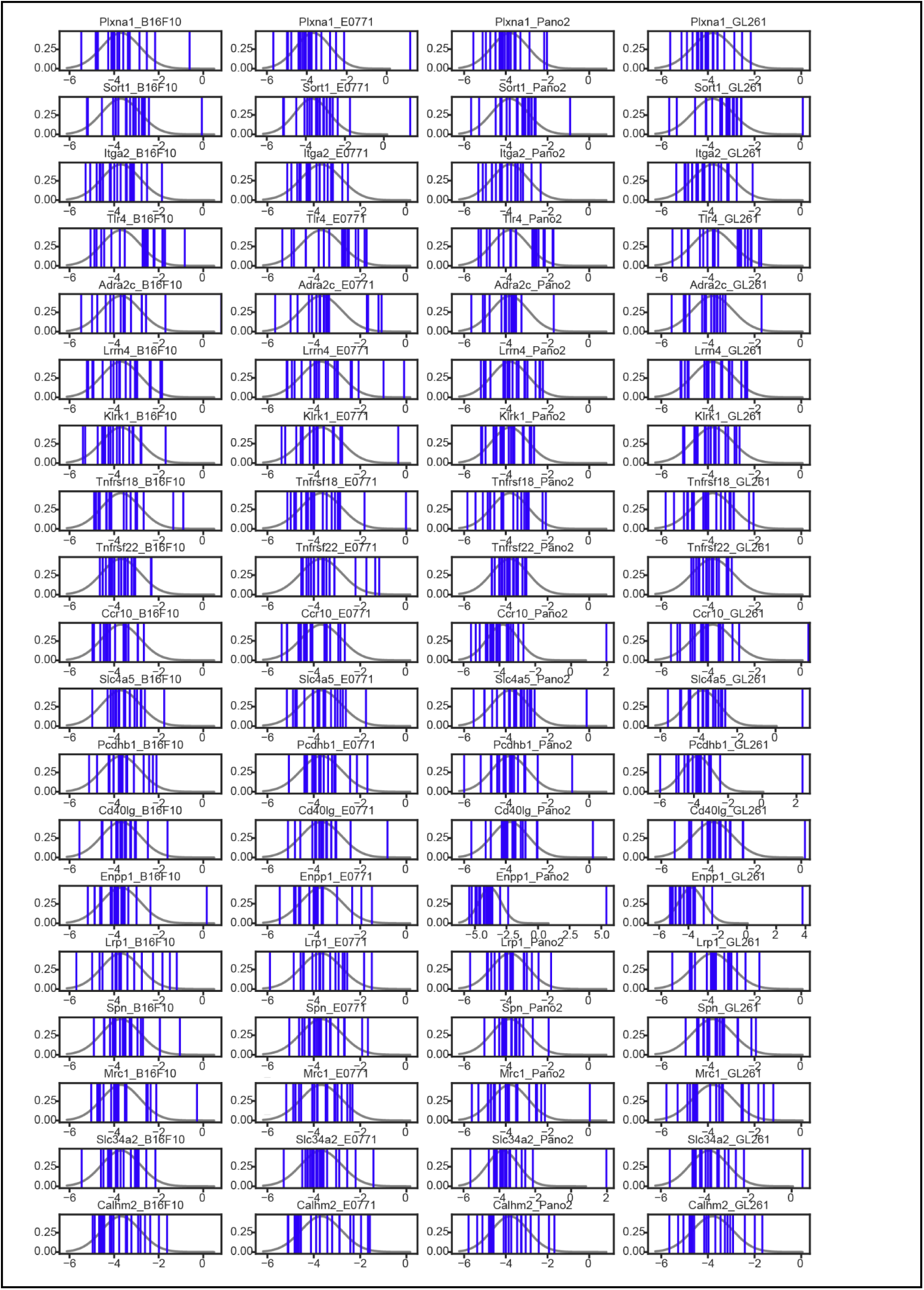
Evaluation of hits reported by external study. The guide-level screening behavior, discretized by tumor model, of genes reported as hits in^[Ref 60]^. The blue lines indicate the log fold changes in guide abundance between natural killer cells before and after injection into tumors. The background distribution (gray) represents the non-targeting negative control guides.

