## Supplementary material for "Balancing off-target and on-target considerations for optimized Cas9 CRISPR knockout library design": Description of Supplementary Data Files.docx.pdf

**Title**: Supplementary Data 1

**Filename**: JacquereGuidePickingProgressions.xlsx

**Description**: A detailed report of the criteria for guide selection in the design of the Jacquere library. For each gene targeted, candidate guides are initially filtered to meet the criteria detailed in picking round 1 and selected in descending order by RS3-seq+target (Chen) on-target score. Criteria are relaxed to those detailed in subsequent picking rounds as necessary to meet the quota of 3 guides per gene. 95.0% of guides in the Jacquere library (quota =3) are selected in picking round 1.

**Title**: Supplementary Data 2

**Filename**: Jacquere_PerGuideAnnotations.csv

**Description**: Guide-level annotations of the Jacquere library with a quota of three guides per gene. Each row reports a unique guide in the Jacquere library along with the gene(s) targeted by the guide. For guides that were selected to target multiple genes, distinct target-level information is delimited by the pipe (" | ") character. The "Other Target Matches" column indicates genes to which each guide sequence maps (in either Ensembl v.113 or Refseq v2024_08) in addition to its intended targets. In the absence of guide mappings to the most current genome annotations, this file is recommended for downstream analysis of Jacquere screening data; please refer to github.com/ldrepano/GPP-Jacquere for example usage. Includes 900 intergenic-targeting and 100 non-targeting guides as negative controls.

**Title**: Supplementary Data 3

**Filename**: Jacquere_PerGuideAnnotations_Quota4.csv

**Description**: Guide-level annotations of the Jacquere library extended to a quota of four guides per gene.

**Title**: Supplementary Data 4

**Filename**: Jacquere_PerTargetAnnotations_uptoquota4.csv

**Description**: The top 4 guide selections per gene targeted in Jacquere to support ad-hoc library design. The "Source" column indicates the source of the gene annotation. If numerous databases recognize the gene, the source is indicated as GENCODE if the gene is part of Ensembl v113 and RefSeq if not. Includes 900 intergenic-targeting and 100 non-targeting guides as negative controls. It is important to note that guides selected to target multiple genes are repeated as an entry for each target gene, and, while the target gene symbols are provided, genes are uniquely identified by their target gene ID.

**Title**: Supplementary Data 5

**Filename**: Jacquere_untargetable_genes.csv

**Description**: Genes not targetable by guides that adhere to Jacquere library design criteria (i.e. CRISPick Aggregate CFD Score <=4.8). "Source" column indicates the gene catalog that recognizes the gene. If numerous catalogs recognize the gene, the source is indicated as GENCODE if the gene is part of Ensembl v113 and RefSeq if not.

**Title**: Supplementary Data 6

**Filename**: Julianna-GRCm38-Ensembl102-CRISPRkoSpCas9-Designs-Quota3.csv

**Description**: Guide-level annotations of the Julianna library with a quota of three guides per gene. The Julianna library implements the same guide selection criteria as Jacquere, only eliminating restrictions on target variability due to the absence of population-level mouse data, against all protein coding genes in the mouse genome build GRCm38 and Ensembl v. 102 annotations. For guides selected to target multiple genes, distinct target-level information is delimited by the pipe (" | ") character. Includes 900 intergenic and 100 non-targeting negative control guides.

**Title**: Supplementary Data 7

**Filename**: Julianna-GRCm38-Ensembl102-CRISPRkoSpCas9-Designs-Quota4.csv

**Description**: Guide-level annotations of the Julianna library extended to a quota of four guides per gene.
